# Bhlhe40 Governs T Cell Effector Differentiation with Distinct Requirements in CD4 and CD8 T Cells for Anti-PD-1 and Anti-CTLA-4 Efficacy

**DOI:** 10.1101/2025.10.10.681497

**Authors:** Akata Saha, Tomoyuki Minowa, Alexander S. Shavkunov, Avery J. Salmon, Sunita Keshari, Suresh Satpati, Nicholas N. Jarjour, Abhinav K. Jain, Kunal Rai, Kristen E. Pauken, Kenneth H. Hu, Brian T. Edelson, Ken Chen, Matthew M. Gubin

**Affiliations:** Department of Immunology, The University of Texas MD Anderson Cancer Center, Houston, TX, USA; Department of Genomic Medicine, The University of Texas MD Anderson Cancer Center, Houston, TX, USA; Department of Pathology and Immunology, Washington University School of Medicine, St. Louis, MO, USA; Department of Epigenetics and Molecular Carcinogenesis, The University of Texas MD Anderson Cancer Center, Houston, TX, USA; James P. Allison Institute, The University of Texas MD Anderson Cancer Center, Houston, TX, USA; Department of Bioinformatics and Computational Biology, The University of Texas MD Anderson Cancer Center, Houston, TX, USA

## Abstract

The transcriptional programs enabling T cells to mediate anti-tumor immunity remain incompletely defined. Here, we identify Bhlhe40 as a key transcriptional regulator that coordinates both CD4 and CD8 T cell effector programs, revealing divergent cell-specific requirements during anti-PD-1 versus anti-CTLA-4 immune checkpoint therapy (ICT). Using conditional knockout mice, we show that anti-PD-1 efficacy depends on CD8 T cell-intrinsic Bhlhe40, whereas anti-CTLA-4 remains effective through Bhlhe40-dependent CD4 T cell Th1 programs that buffer impaired effector function in Bhlhe40-deficient CD8 T cells. Mechanistically, loss of Bhlhe40 reduces IFN-γ production and skews CD8 T cells toward TCF-1-expressing progenitor exhausted/stem-like states at the expense of effector differentiation, impairing glycolytic fitness under both therapies and mitochondrial function during anti-PD-1 treatment, thereby revealing a Bhlhe40-dependent coupling between effector differentiation, cytokine production, and metabolic fitness that is particularly critical for anti-PD-1 efficacy. CD8 T cell-intrinsic Bhlhe40 also promotes critical ICT-induced remodeling from M2-like CX3CR1 macrophages to inflammatory iNOS macrophages. Analysis of human cancer datasets supported our preclinical observations, revealing that *BHLHE40* is enriched in tumor-reactive and activated/exhausted CD8 T cells, where its expression is inversely correlated with *TCF7* (TCF-1) and positively associated with *TOX, GZMB,* and *IFNG*. Moreover, persistent CD8 T cell clones from basal cell carcinoma responders exhibited significantly higher *BHLHE40* expression at pre-treatment than those from non-responders to PD-1 blockade. Together, these findings establish Bhlhe40 not only as a transcriptional coordinator of T cell effector programs, but also as a therapy-specific, subset-dependent determinant that differentially governs CD4 and CD8 T cell contributions to anti-PD-1 and anti-CTLA-4 efficacy.

## INTRODUCTION

Immune checkpoint therapy (ICT) targeting PD-1 or CTLA-4 can elicit durable tumor regression in some patients, but many fail to respond^1–7^. While tumor-reactive T cells are central to ICT efficacy^8–16^, the transcriptional programs that enable CD4 and CD8 T cells to mediate effective anti-tumor immunity remain incompletely defined^17–19^. Understanding these requirements is important because anti-PD-1 and anti-CTLA-4 each act through both overlapping as well as distinct mechanisms^20,21^. Basic helix-loop-helix family member e40 (Bhlhe40) has emerged as a critical transcription factor regulating immune responses^22^. Bhlhe40 often functions as a transcriptional repressor, which in some contexts involves binding promoter or enhancer regions of target loci^22–24^; however, it can also function as a transcriptional activator, particularly in cooperation with other transcription factors^25^. Bhlhe40 has also been shown to indirectly enhance gene transcription by sustaining intracellular acetyl-CoA to support histone acetylation^26^. Bhlhe40 is expressed across multiple cell types^27,28^, including T cells, where its expression is driven by TCR engagement and CD28 co-stimulation, can be induced by inflammatory cytokines such as GM-CSF and IL-1β, and is restrained by PD-1 signaling^22,26,29,30^. Mice lacking Bhlhe40 in both CD4 and CD8 T cells are resistant to experimental autoimmune encephalomyelitis, reflecting the requirement for Bhlhe40 to repress IL-10 and promote GM-CSF production in T cells^23,27,31^. Conversely, these mice exhibit increased susceptibility to certain infections^24,32^.

We recently demonstrated neoantigen (NeoAg)-specific intratumoral CD4 and CD8 T cells upregulate Bhlhe40 in tumor-bearing mice treated with anti-PD-1 or anti-CTLA-4 ICT^30^. Using CD4-Cre Bhlhe40^fl/fl^ mice—where deletion occurs in both CD4 and CD8 T cells—we found that T cell-intrinsic Bhlhe40 is essential for tumor rejection in response to either ICT. Global deletion of Bhlhe40 disrupted T cell effector gene expression and impaired ICT-driven remodeling of the tumor microenvironment (TME). Independent work demonstrated Bhlhe40 is required for response to anti-PD-1/PD-L1 ICT in preclinical models, with loss of T cell-intrinsic Bhlhe40 linked to defective mitochondrial fitness^26^. Thus, a central unresolved question is whether Bhlhe40 functions as a shared transcriptional node across immunotherapies or instead plays distinct, T cell-subset-specific roles that differentially govern responses to anti-PD-1 versus anti-CTLA-4, and through what mechanisms these effects are mediated.

Here, we dissect the cell-type-specific requirements for Bhlhe40 in ICT by selectively deleting it in CD8 T cells or in both CD4 and CD8 T cells using tumor models where both subsets are essential for therapeutic response. We show that CD8 T cell-intrinsic Bhlhe40 is required for tumor control by anti-PD-1, with additional contributions from CD4 T cell-intrinsic Bhlhe40, whereas anti-CTLA-4 efficacy is largely dependent on CD4 T cell-intrinsic Bhlhe40 and remains intact when Bhlhe40 is absent from CD8 T cells. Loss of Bhlhe40 skews CD8 T cells toward a more naïve or progenitor exhausted-like state and reduced acquisition of terminal effector features, particularly in the absence of ICT. In addition, Bhlhe40 supports glycolysis under both anti-PD-1 and anti-CTLA-4 therapies, sustains mitochondrial respiration primarily during anti-PD-1, and is essential for full ICT-induced myeloid remodeling from CX3CR1 to iNOS macrophages. Analysis of multiple human cancer datasets demonstrates that *BHLHE40* is enriched in tumor-reactive and activated/exhausted CD8 T cell populations across tumor types, inversely correlated with *TCF7*, and positively associated with *TOX, GZMB,* and *IFNG*. Moreover, persistent CD8 T cell clones from basal cell carcinoma responders exhibited higher *BHLHE40* expression at pre-treatment than those from non-responders to PD-1 blockade. Our findings establish Bhlhe40 as a key regulator of T cell effector programming and as a transcriptional determinant of subset-specific functions that underlie response to anti-PD-1 versus anti-CTLA-4 therapy.

## RESULTS

### Human Tumor-Reactive CD4 and CD8 Effector T Cells Express Bhlhe40

We previously found that Bhlhe40 is expressed in intratumoral CD4 and CD8 T cells and mice lacking Bhlhe40 in both CD4 and CD8 T cells fail to respond to anti-PD-1 or anti-CTLA-4 immune checkpoint therapy (ICT)^30^. To extend these findings to human tumors, we first examined *BHLHE40* expression across immune cell types inferred from The Cancer Genome Atlas (TCGA) RNA-sequencing (RNA-seq) data. Both CD4 and CD8 T cells consistently expressed *BHLHE40* across multiple cancer types (**Fig. 1A**). We next asked whether its expression is selectively enriched within human tumor-reactive CD8 T cells, since tumors can contain both tumor-specific CD8 T cells as well as non-tumor-reactive bystander CD8 T cells^33^. Tumor-reactive CD8 T cells were identified in studies by Oliveira et al. and Minowa et al.^34,35^, which analyzed human cutaneous melanoma (an immunogenic tumor type that frequently responds to ICT), and acral melanoma (a less immunogenic tumor type), respectively. Reanalysis of datasets from both these studies revealed enrichment of *BHLHE40* in tumor-reactive CD8 T cells compared with predominately non-tumor-reactive CD8 T cells (**Fig. 1B**). Overall, these findings align with our observations in murine models, where Bhlhe40 is highly expressed within tumor-reactive populations^30^.

**Figure 1.**
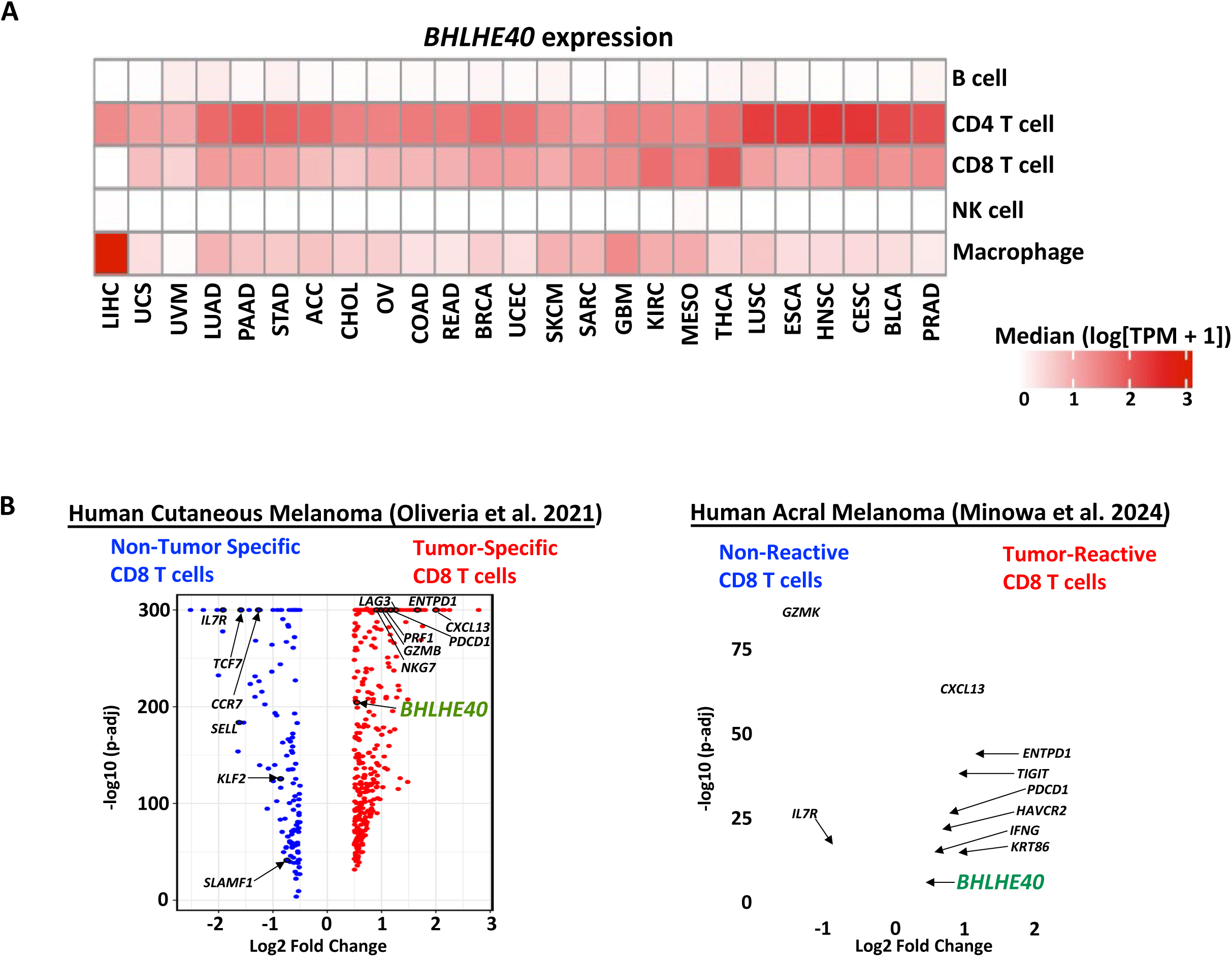
Bhlhe40 is expressed in tumor-reactive human T cells. **(A)** *BHLHE40* transcript levels across immune cell types and various tumor types from The Cancer Genome Atlas (TCGA) datasets analyzed via GEPIA2021 (https://doi.org/10.1093/nar/gkab418). **(B)** Differential gene expression analysis comparing tumor-reactive CD8 T cells with non-tumor-specific or non-reactive CD8 T cells (as defined by Oliveira et al. and Minowa et al.) in cutaneous and acral melanoma.

### CD8 T Cell-Intrinsic Bhlhe40 Is Essential for Anti-PD-1 but Not Anti-CTLA-4 Efficacy

We previously reported that *CD4-Cre Bhlhe40^fl/fl^* (B^ΔT^) conditional knockout (KO) mice lacking Bhlhe40 in both CD4 and CD8 T cells (because CD4-Cre is expressed in double-positive thymocytes^36^) fail to respond to anti-PD-1 or anti-CTLA-4 ICT^30^. Consequently, we first sought to determine whether Bhlhe40 expression specifically within CD8 T cells is required for the therapeutic efficacy of anti-PD-1 and anti-CTLA-4. We used *CD8 E8I-Cre* mice, in which Cre recombinase is driven by the E8I CD8 enhancer that is active exclusively in CD8 T cells^37^. We verified Bhlhe40 was specifically deleted in intratumoral CD8 T cells (and not CD4 T cells) in isotype control antibody (Ab)-, anti-PD-1-, or anti-CTLA-4- treated *CD8 E8I-Cre Bhlhe40^fl/fl^* (B^ΔCD8^) mice bearing the 1956 MCA sarcoma tumor^30,38,39^ (**Fig. S1A**). Using this well-characterized tumor model, we next injected 1956 tumor cells into Bhlhe40-sufficient *Bhlhe40^fl/fl^* (B^f/f^) control mice, B^ΔT^ mice (Bhlhe40 KO in CD4 and CD8 T cells), B^ΔCD8^ mice (Bhlhe40 KO in CD8 T cells) and treated them with control Ab, anti-PD-1, or anti-CTLA-4 (**Fig. 2A**). Consistent with our prior findings^30^, tumor outgrowth was observed in B^ΔT^ mice treated with anti-PD-1 or anti-CTLA-4 (**Fig. S1B**). While tumor regression occurred in anti-PD-1-treated B^f/f^ control mice, anti-PD-1 failed to control tumor growth in B^ΔCD8^ mice (**Fig. 2B**). Interestingly, anti-CTLA-4 elicited effective tumor control in both B^f/f^ and B^ΔCD8^ mice (**Fig. 2B**). We obtained similar results using the *Braf^V600E^ Pten^−/−^ Cdkn2^−/−^* YUMM1.7^40^-modified Y1.7LI melanoma model^41^ (**Figs. S1C and S1D**). To confirm that expression of Cre itself does not affect anti-tumor immune responses, we compared *CD8 E8I-Cre Bhlhe40^+/+^* mice to B^f/f^ mice and observed comparable tumor growth and responses to ICT (**Fig. S1E**). Altogether, these results demonstrate that CD8 T cell-intrinsic Bhlhe40 is required for anti-PD-1 efficacy but is dispensable for response to anti-CTLA-4.

**Figure 2.**
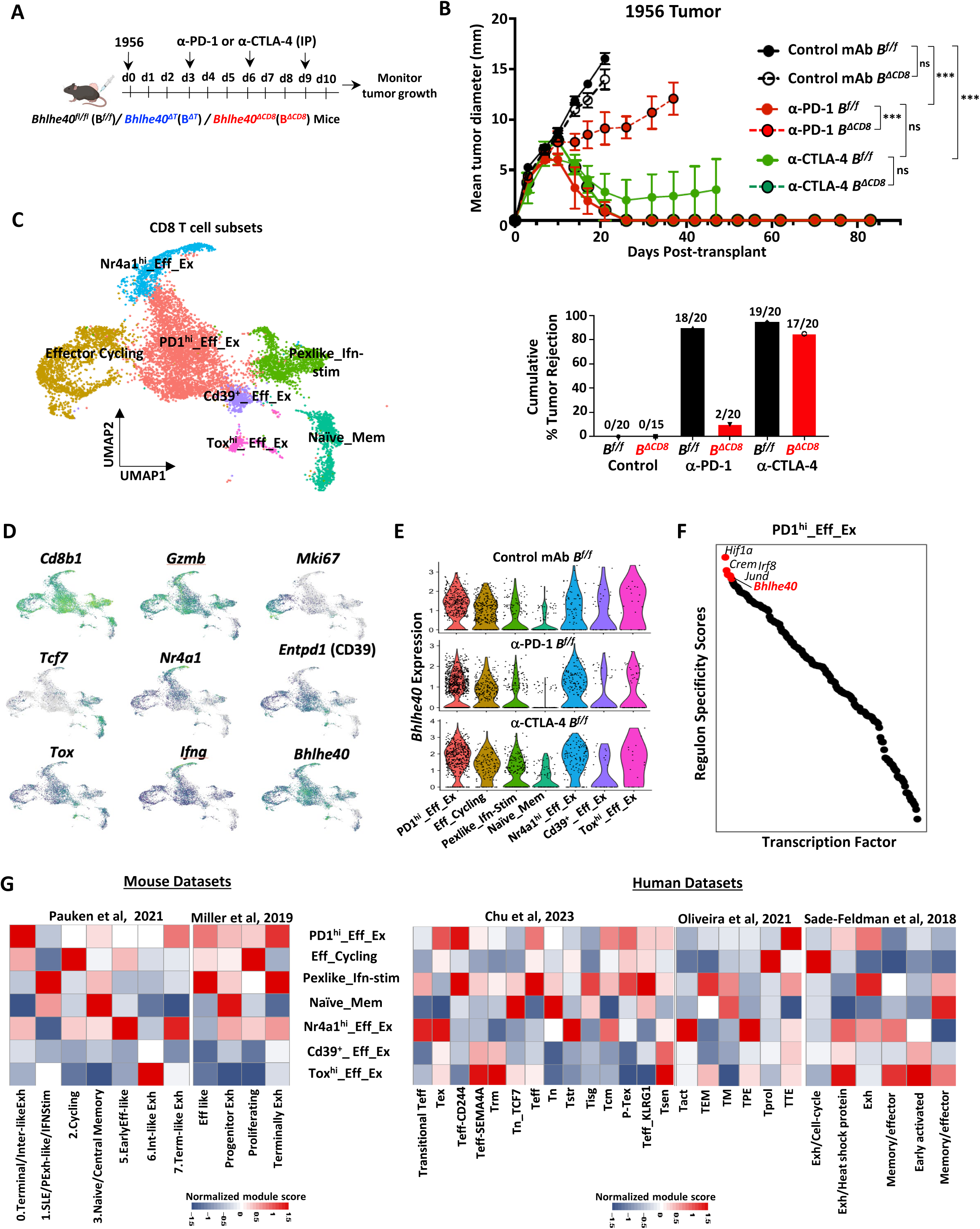
CD8 T cell-intrinsic Bhlhe40 is essential for anti-PD-1 but not anti-CTLA-4 efficacy. **(A)** Bhlhe40 conditional knockout mice bearing 1956 MCA sarcoma were treated with anti-PD-1 or anti-CTLA-4 as indicated. **(B)** Tumor growth and percent tumor rejection of 1956 tumors in Bhlhe40^fl/fl^ (B^f/f^) and Bhlhe40^ΔCD8^ (B^ΔCD8^) mice treated with control antibody (Ab), anti-PD-1, or anti-CTLA-4. **(C)** UMAP generated from integration of CD8 T cells from scRNAseq Experiment #1 and scRNAseq Experiment #2 depicting CD8 T cell clusters named based on phenotypic markers. **(D)** Feature plots displaying expression of select markers. **(E)** Violin plots showing *Bhlhe40* expression in CD8 T cells by cluster and treatment in B^f/f^ mice. **(F)** Dot plot showing the regulon specificity score across transcription factors. The top five regulons and Bhlhe40 in PD1^hi^_Eff_Ex cluster are highlighted in red and labeled on the plot. **(G)** Heatmaps comparing features (module scores) of CD8 T cell clusters (rows) to published mouse and human CD8 T cell gene signatures (columns) identified/annotated by Miller et al. and Pauken et al. (mouse) and by Chu et al., Oliveira et al., and Sade-Feldman et al. (human). For (**B**), tumor growth is presented as mean +/− SEM of mean tumor diameter of individual mice and are representative of at least three independent experiments (∗∗∗p < 0.001; ns, not significant, 2-way ANOVA). Tumor rejection graphs display cumulative percentage of mice with complete tumor rejection from 3-4 independent experiments.

### Bhlhe40 Is Enriched in Effector/Exhausted CD8 T Cells

To gain insights into the divergent effects of anti-PD-1 and anti-CTLA-4 therapies, we performed single-cell RNA sequencing (scRNAseq) on immune cells isolated from 1956 tumors of Bhlhe40-sufficient (B^f/f^) mice, mice lacking Bhlhe40 only in CD8 T cells (B^ΔCD8^), and mice lacking Bhlhe40 in both CD4 and CD8 T cells (B^ΔT^) treated with control Ab, anti-PD-1, or anti-CTLA-4 (**Fig. S2A**). Live Thy1.2^+^ CD45^+^ and Thy1.2^−^ CD45^+^ immune cells were sorted, barcoded, and mixed at 60:40 ratio to enhance capture of less frequent lymphoid cells for scRNAseq. To further increase lymphocyte resolution by accumulating more lymphocytes, we performed an additional experiment under the same treatment conditions, this time focusing specifically on Thy1.2 CD45 lymphocytes from B^f/f^ and B^ΔCD8^ mice (**Fig. S2A**). We then integrated the scRNAseq data from both experiments for downstream analysis of CD4 and CD8 T cells (**Fig. S2B**).

Unsupervised clustering of *Cd3e^+^ Cd8a^+^* cells from our scRNAseq experiment identified seven CD8 T cell clusters (plus two small clusters of non-CD8 T cells, which were removed from downstream analyses). These spanned a range of states and included activated effector or exhausted (PD1^hi^_Eff_Ex; Nr4a1^hi^_Eff_Ex; Cd39^+^ Eff_Ex; Tox^hi^_Eff_Ex), cycling/proliferating cells (Eff_Cycling), progenitor exhausted (Tpex)-like/interferon-stimulated cells (Tpexlike_Ifn-Stim), and naïve/memory-like populations (Naïve_Mem) based on functional marker transcript expression and comparisons to published datasets^34,42–45^ (**Figs. 2C-2G, S2C,** and **S3**).

Treatment with ICT reshaped the CD8 T cell compartment in a manner that was dependent on genotype and whether mice were treated with anti-PD-1 or anti-CTLA-4 **(Fig. S2E)**. We first noted that effector/exhausted clusters expressed the highest levels of *Bhlhe40* in B^f/f^ mice. Among these, the PD1^hi^_Eff_Ex cluster exhibited the highest inferred Bhlhe40 regulon activity by SCENIC analysis, whereas the Naïve_mem cluster showed the lowest regulon activity (**Figs. 2D-F, S2C,** and **S2D**). PD1^hi^_Eff_Ex also expressed key effector molecules (e.g., *Ifng*, *Gzmb*, *Prf1*), as well as relatively high levels of inhibitory receptor transcripts (e.g., *Pdcd1* (PD-1), *Havcr2* (TIM-3), *Lag3*), which are induced upon TCR activation but become hallmarks of exhausted and/or dysfunctional CD8 T cells when their expression is high and sustained^46–50^ (**Figs. S2C** and **S3**). Nr4a1^hi^_Eff_Ex was noted for its expression of *Nr4a1* (Nur77), a gene upregulated proportionally to TCR signaling strength^51,52^, as well as high levels of *Ifng, Gzmb,* and *Cd69* (**Figs. 2D**, **S2C**, and **S3**). Tox^hi^_Eff_Ex was the smallest cluster and expressed high levels of *Tox,* encoding a transcription factor essential for enforcing exhausted/dysfunctional T cell states^53–56^, in addition to *Pdcd1, Havcr2*, *Lag3*, and *Entpd1* (CD39) (**Figs. 2D**, **S2C**, and **S3**). Cells within this cluster also expressed *Slamf6* (Ly108) and *Tcf7*, which encodes the TCF-1 transcription factor that supports stem-like Tpex cells essential for sustained anti-PD-1 responses and for anti-CTLA-4-driven T cell memory^17,43,57–62^. Notably, expression of *Tcf7* was restricted to a subset of cells and was partially exclusive of *Tox* expression (**Figs. 2D** and **S2F**). Clusters Eff_Cycling was a proliferating cluster that expressed *Bhlhe40* at moderate levels, whereas Tpex-like_Ifn-stim expressed lower levels of *Bhlhe40* (**Fig. S2C**). Cluster Tpex-like_Ifn-stim displayed upregulation of interferon (IFN)-stimulated transcripts, high *Gzmk* expression, and also Tpex-associated transcripts such *Tcf7* and *Slamf6* **(Figs. S2C** and **S3**).

### Bhlhe40 Regulates Neoantigen-Specific CD8 T Cell Responses to Anti-PD-1 and Supports IFN-**γ** Production

We assessed whether the differential tumor control observed with anti-PD-1 versus anti-CTLA-4 in the absence of CD8 T cell-intrinsic Bhlhe40 was reflected at the level of tumor-specific T cell responses. Anti-CTLA-4 increased the overall frequency of CD8 T cells in B^f/f^ mice, while no significant changes were observed across other conditions (**Fig. 3A**). However, both anti-PD-1 and anti-CTLA-4 increased the frequency of tumor-specific CD8 T cells recognizing mutant Psmd6, an MHC class I (MHC-I) mutant NeoAg in the 1956 tumor^38^ (**Fig. 3B**). Notably, anti-CTLA-4 drove this increase independently of CD8 T cell-intrinsic Bhlhe40, whereas anti-PD-1 selectively increased Psmd6-specific CD8 T cells in B^f/f^ control mice and not in B^ΔCD8^ mice.

**Figure 3.**
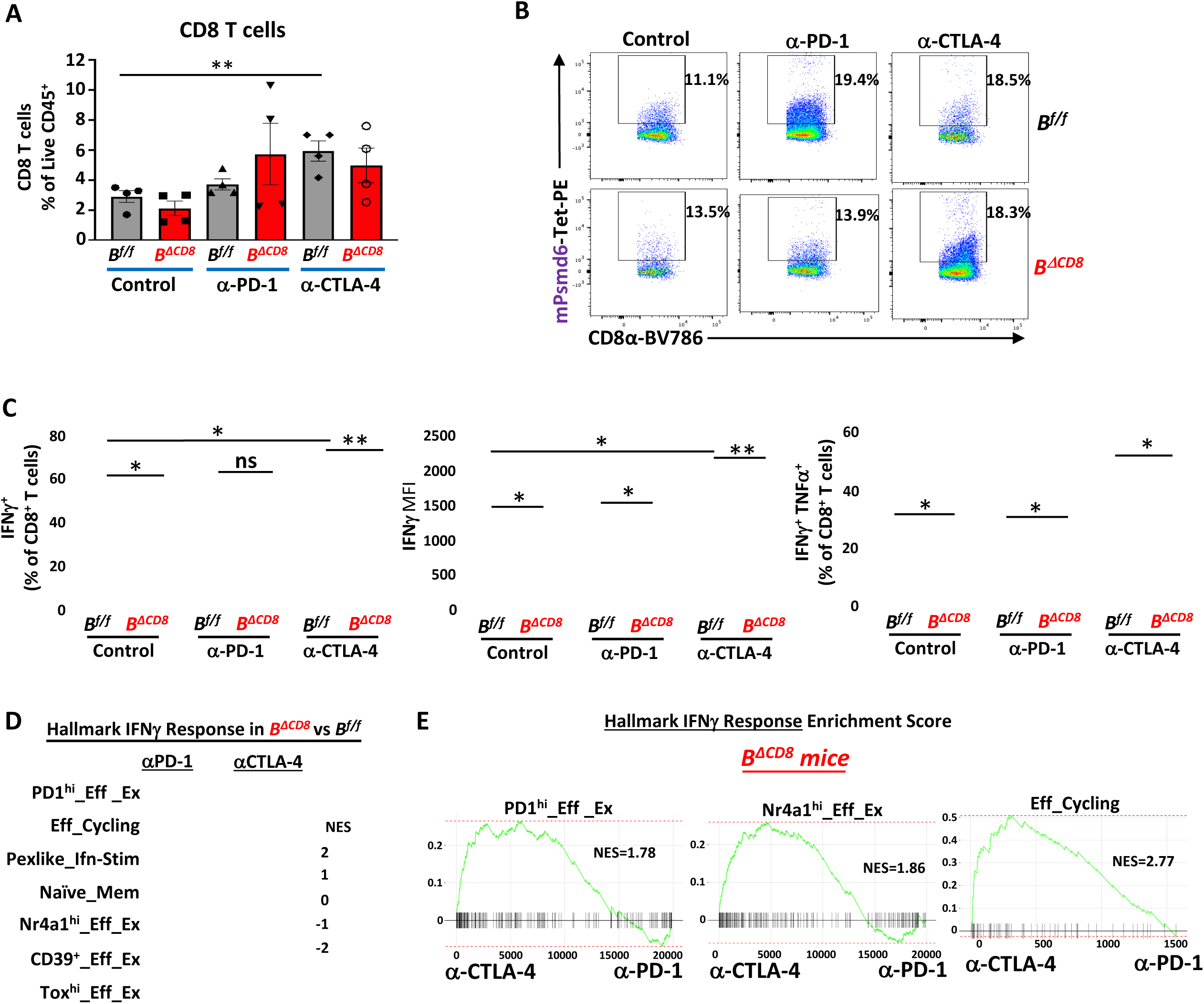
Bhlhe40 regulates NeoAg-specific CD8 T cell responses to anti-PD-1 and supports IFN-γ production. **(A)** Bar graph showing percent of CD8 T cells as determined by flow cytometry staining of intratumoral cells isolated from 1956 tumor-bearing B^f/f^ and B^ΔCD8^ mice treated with control Ab, anti-PD-1, or anti-CTLA-4. **(B)** Flow cytometry plots of mPsmd6 peptide-H2-K^b^ tetramer staining for NeoAg-specific CD8 T cells in B^f/f^ and B^ΔCD8^ mice treated with control Ab, anti-PD-1, or anti-CTLA-4. Gated on live CD8 T cells. **(C)** Graphs displaying IFN-γ^+^CD8 T cells (as a percentage of CD8 T cells), IFN-γ MFI of IFN-γ^+^CD8 T cells, and IFN-γ^+^ TNF-α^+^CD8 T cells (as a percentage of CD8 T cells). **(D)** Heatmap displaying normalized Hallmark IFNγ response GSEA enrichment score (NES) for CD8 T cell clusters in *B*^Δ*CD*8^ versus *B^f/f^* mice treated with anti-PD-1 or anti-CTLA-4. **(E)** GSEA plots for the Hallmark IFNγ response in PD1^hi^_Eff_Ex, Nr4a1^hi^_Eff_Ex and Eff_Cycling CD8 T cell clusters in B^ΔCD8^ mice comparing anti-CTLA-4 vs anti-PD-1 treatment. For Bar graphs in (**A**) display the mean ± SEM and are representative of at least three independent experiments (∗∗p < 0.01; ns, not significant, unpaired t test). For (**C**), IFN-γ and TNF-α was determined via intracellular cytokine staining of *ex vivo* restimulated CD8 T cells isolated from 1956 tumors on day 10 post-tumor transplant. Bar graphs display the mean ± SEM and are representative of at least three independent experiments (∗∗p < 0.01, ∗∗∗p < 0.001; ns, not significant, unpaired t test).

We previously found that CD8 T cells from mice lacking Bhlhe40 in both CD4 and CD8 T cells exhibited altered responses to ICT, including defects in IFN-γ expression^30^; however, it was unclear whether these changes were due to a CD8 T cell-intrinsic requirement for Bhlhe40 or reflected indirect effects stemming from its ablation in CD4 T cells. Our use of B^ΔCD8^ mice now demonstrate that some of these Bhlhe40-dependent changes, including reduced IFN-γ production, are at least partially attributable to CD8 T cell-intrinsic Bhlhe40. Specifically, cluster Nr4a1^hi^_Eff_Ex, which exhibited the highest levels of *Ifng* among CD8 T cell clusters (**Figs. S2C** and **S3**), as well as several other clusters showed reduced *Ifng* expression in B^ΔCD8^ mice compared to B^f/f^ mice within the corresponding treatment groups (**Fig. S4A**). We observed an even more pronounced reduction of *Ifng* expression in mice lacking Bhlhe40 in both CD8 and CD4 T cells (B^ΔT^), suggesting that Bhlhe40 expression in CD4 T cells also supports *Ifng* production by CD8 T cells (**Fig. S4A**). At the protein level, although IFN-γ-positive cells as a percentage of CD8 T cells did not change significantly between B^f/f^ and B^ΔCD8^ mice with anti-PD-1 treatment, the expression level of IFN-γ was significantly decreased based on mean fluorescence intensity (MFI) (**Fig. 3C**). Notably, reductions in IFN-γ production were observed not only with anti-PD-1—which fails to induce rejection in B^ΔCD8^ mice—but also with anti-CTLA-4, despite the latter still leading to tumor rejection in B^ΔCD8^ mice. Further, a significant decrease in IFN-γ^+^ TNF-α^+^ CD8 T cells was observed in B^ΔCD8^ mice compared to B^f/f^ mice within comparable treatment groups (**Fig. 3C**). In addition to alterations in *Ifng*, *Il10* expression was increased in the clusters expressing the highest levels of *Il10* (PD1^hi^_Eff_Ex and Nr4a1^hi^_Eff_Ex) in B^ΔCD8^ mice (**Fig. S4B**), consistent with Bhlhe40’s function as a transcriptional repressor of *Il10*^23,24,28^.

Gene set enrichment analysis (GSEA) revealed that, under anti-PD-1 or anti-CTLA-4 treatment, multiple CD8 T cell clusters from B^ΔCD8^ mice exhibited significant negative enrichment of the Hallmark IFN-γ response gene set compared to B^f/f^ controls, with several clusters also showing reduced IFN-α response signatures (**Figs. 3D**, **S5**, and **S6**). Reduced IFN-γ signatures likely reflect impaired IFN-γ production by Bhlhe40-deficient CD8 T cells, whereas diminished IFN-α response may represent secondary effects of altered CD8 T cell function. Notably, although IFN-γ response genes were reduced in B^ΔCD8^ mice under anti-CTLA-4 relative to B^f/f^ controls, they remained relatively enriched compared to anti-PD-1-treated B^ΔCD8^ mice (**Fig. 3E**).

### Bhlhe40 Shapes Effector and Exhausted CD8 T Cell States

We hypothesized that Bhlhe40 contributes to effector/exhausted phenotypes as we noted in several clusters, expression of not only *Ifng*, but also *Pdcd1, Lag3, Ctla4, Gzmk,* and the fractalkine receptor gene *Cx3cr1* were reduced in B^ΔCD8^ mice relative to B^f/f^ controls (**Figs. S3** and **S4A**). These cells also showed increased expression of several immunoregulatory genes (*Cd5*, *Cd6*, *Ptprj*, and *Pag1*) (**Figs. S3** and **S4A**). We also observed changes that followed similar patterns in control Ab and anti-PD-1 conditions but diverged under anti-CTLA-4 treatment. Under anti-PD-1 treatment, induction of *Prf1* was attenuated in B^ΔCD8^ mice, which also showed lower baseline *Prf1* levels under control treatment, whereas anti-CTLA-4 treatment resulted in higher *Prf1* expression in B^ΔCD8^ compared to B^f/f^ mice (**Figs. S3** and **S4A**). Similarly, *Tox* expression was reduced in anti-PD-1-treated B^ΔCD8^ mice, whereas anti-CTLA-4 reduced *Tox* expression to comparable levels in both B^f/f^ and B^ΔCD8^ mice (**Fig. S4C**). A notable exception to the reduced expression of effector and exhaustion-associated genes was increased *Gzmb* expression in Bhlhe40-deficient CD8 T cells upon anti-PD-1 or anti-CTLA-4 treatment.

To better understand how Bhlhe40 shapes these phenotypes, we explored its relationship with CD8 T cell phenotype-defining transcriptional programs. Correlation analysis revealed an inverse relationship between *Bhlhe40* and *Tcf7*, *Sell*, *Id3*, *Bcl2*, *Cxcr3*, *Lef1*, *Slamf6*, and *Eomes*, genes associated with naïve, memory, and Tpex-like phenotypes (**Figs. 4A** and **S7A**). In contrast, *Bhlhe40* was associated with transcripts linked to T cell activation, survival, effector differentiation, and exhaustion including *Nr4a1, Junb, Jund, Bcl2a1b, Rgs16, Irf8, Rbpj,* and *Dusp5*^63–65^ as well as inhibitory receptors (*Pdcd1*, *Ctla4, Lag3, Havcr2, Klrc1* [NKG2A]). Bhlhe40 was also correlated with genes related to cytotoxicity (*Nkg7*, *Srgn*) and costimulatory receptors (*Tnfrsf4* (OX40), *Tnfrsf9* (4-1BB), *Icos*), and *Hif1a*, a TCR-induced transcript whose stability is enhanced under hypoxic conditions, together with the downstream HIF1-responsive gene *Hilpda* (**Figs. 4A** and **S7A**).

**Figure 4.**
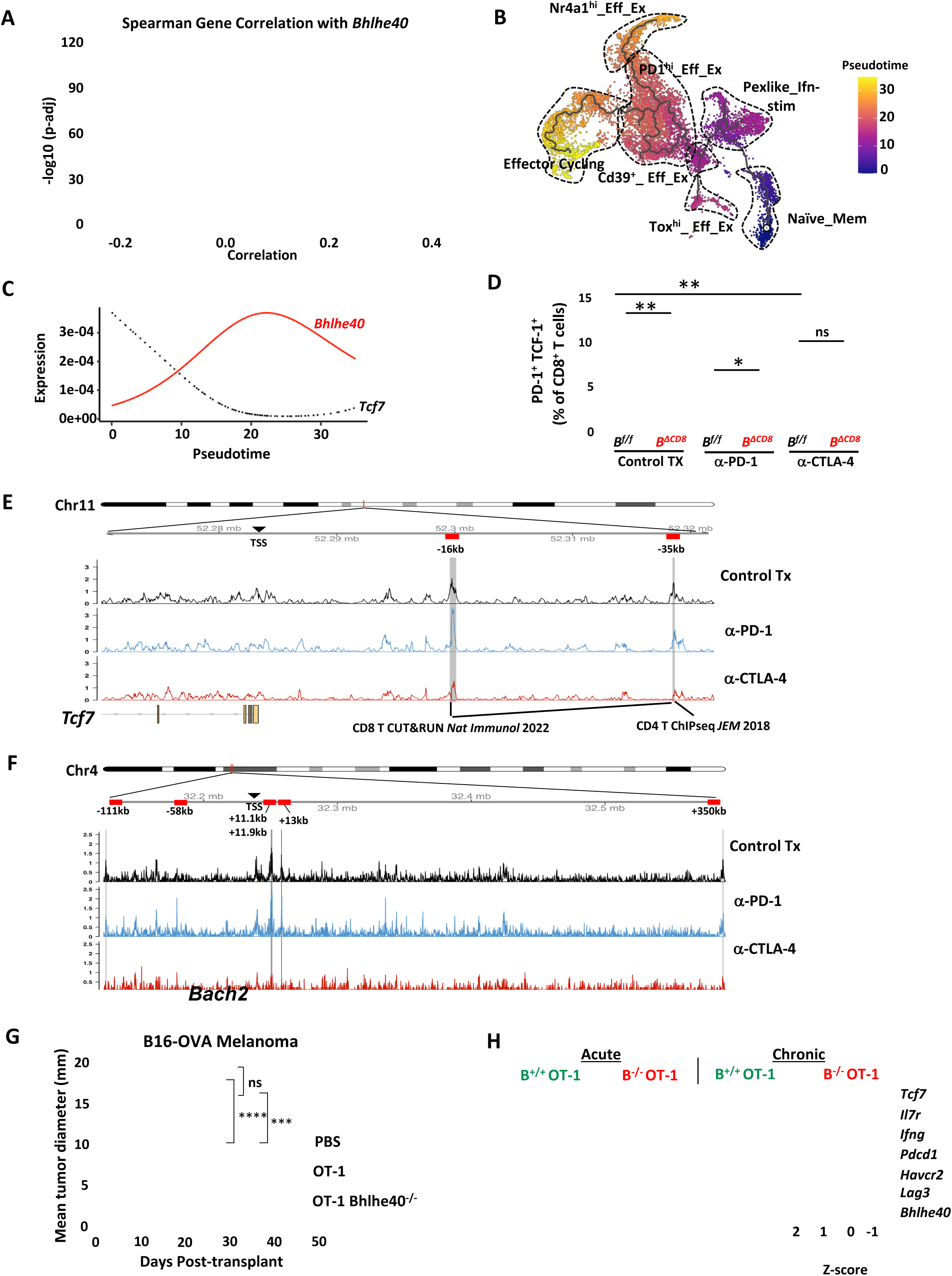
CD8 T cell-intrinsic Bhlhe40 governs effector differentiation. **(A)** Spearman correlation analysis of all CD8 T cell clusters relative to Bhlhe40 compared in B^f/f^ mice. Dot color indicates the p-value. **(B)** UMAP of CD8 T cells showing the Monocle3 trajectory across all samples, colored by inferred pseudotime. **(C)** Pseudo-temporal trends of *Bhlhe40* and *Tcf7* in CD8 T cells from B^f/f^ mice. **(D)** Graphs displaying PD-1^+^TCF-1^+^ CD8 T cells (as a percentage of CD8 T cells). **(E)** Genome track plots showing Bhlhe40 CUT&RUN peaks at *Tcf7* locus in intratumoral CD8 T cells isolated from WT mice treated with control Ab, anti-PD-1, or anti-CTLA-4. **(F)** Genome track plots showing Bhlhe40 CUT&RUN peaks at *Bach2* locus in intratumoral CD8 T cells isolated from WT mice treated with control Ab, anti-PD-1, or anti-CTLA-4. **(G)** Tumor growth of B16-OVA tumors in mice adoptively transferred with either PBS, Bhlhe40^+/+^ (B^+/+^) OT-I or Bhlhe40^-/-^ (B^-/-^) OT-I splenocytes stimulated ex vivo with SIINFEKL peptide. **(H)** Heatmap displaying transcript expression of effector/exhaustion markers (*Pdcd1, Lag3, Havcr2, Ifng*) and memory-associated markers (*Tcf7, Il7r*) in B^+/+^ and B^-/-^ OT-I T cells under ex vivo acute and chronic stimulation conditions with cognate SIINFEKL peptide-loaded APCs. For (**D**), TCF-1 and PD-1 expression was assessed via intracellular/intranuclear staining and flow cytometry of 1956 tumors on day 10 post-tumor transplant. Bar graphs display the mean ± SEM and are representative of two independent experiments (∗p < 0.05; ∗∗p < 0.01, ns, not significant, unpaired t test). Tumor growth in (**G**) is presented as mean +/− SEM of mean tumor diameter of individual mice and is representative of two independent experiments (∗∗∗p < 0.001; ∗∗∗∗p < 0.0001; ns, not significant, 2-way ANOVA).

Given the strong inverse correlation between *Bhlhe40* and *Tcf7*-associated naïve/memory and Tpex programs, we next examined how loss of Bhlhe40 affected these states. The Naïve_mem cluster expressed the highest levels of *Tcf7* (**Figs. S3, S8A**, and **S8B**), and this cluster was markedly expanded in control Ab-treated B^ΔCD8^ mice compared to B^f/f^ mice (**Fig. S2E**). Under anti-PD-1 treatment, *Tcf7* expression within the Naïve_mem cluster was also elevated in B^ΔCD8^ mice. Moreover, in the absence of ICT, increased *Tcf7* expression in B^ΔCD8^ mice was observed across several CD8 T cell clusters in which only a small fraction of cells expressed *Tcf7*. A similar pattern was noted for *Bach2*, which encodes a transcriptional repressor that enforces stem-like and Tpex states^66^ (**Fig. S8B**). Consistent with these findings, the Naïve_mem cluster showed the lowest putative Bhlhe40 regulon activity (**Fig. S2D**). Trajectory analysis further supported this relationship, as Monocle3 analysis starting from Naïve_mem revealed opposing dynamics of *Bhlhe40* and *Tcf7* across pseudotime (**Figs. 4B** and **4C**). At the protein level, intratumoral Tpex/Tpex-like PD-1 TCF-1 CD8 T cells were significantly expanded in B^ΔCD8^ mice under both control Ab and anti-PD-1 treatment conditions compared to B^f/f^ mice (**Fig. 4D**).

We next sought to determine whether Bhlhe40 directly interacts with regulatory regions of genes associated with these states by performing CUT&RUN to map Bhlhe40 binding sites in intratumoral CD8 T cells from WT mice treated with control Ab, anti-PD-1, or anti-CTLA-4. Consistent with this hypothesis, Bhlhe40 occupied distal intergenic regions of *Tcf7* under control conditions (−16 kb and −35 kb from the transcription start site [TSS]) and under anti-PD-1 treatment (−35 kb from the TSS) (**Fig. 4E** and **Table S6-S7**). Comparison with published datasets revealed that the −16 kb and −35 kb regions are consistent with Bhlhe40 CUT&RUN peaks in CD8 T cells from LCMV-infected mice^67^, whereas the −35 kb region also overlaps a Bhlhe40 ChIP-seq peak identified in *in vitro* polarized CD4 T cells^24^. We also identified five Bhlhe40 CUT&RUN peaks within the *Bach2* locus under control treatment conditions and four under anti-PD-1 treatment (**Fig. 4F** and **Table S6-S7**). Together, these findings support a model in which Bhlhe40 directly restrains *Tcf7*- and *Bach2*-associated Tpex-like programs while promoting effector differentiation in tumor-infiltrating CD8 T cells.

### Bhlhe40 Is Required for Differentiation Toward Conventional Effector and Exhausted CD8 T Cell States

To further examine how Bhlhe40 loss influences the balance between effector/exhausted and Tpex-like CD8 T cell phenotypes, we crossed OT-I TCR transgenic mice with Bhlhe40^-/-^ mice to generate Bhlhe40-deficient OT-I (B^-/-^ OT-I) mice. To ensure *in vivo* relevance of this system, we first evaluated whether Bhlhe40 was required for OT-I CD8 T cell-mediated tumor control in an adoptive cell therapy model. Splenocytes from B^+/+^ OT-I and B^-/-^ OT-I mice were stimulated with SIINFEKL peptide, the ovalbumin (OVA)-derived cognate epitope presented by H2-K^b^. Stimulated B^+/+^ or B^-/-^ OT-I cells were then isolated and adoptively transferred into mice bearing B16-OVA melanomas. While transfer of B^+/+^ OT-I CD8 T cells delayed tumor outgrowth, B^-/-^ OT-I CD8 T cells failed to reduce tumor growth (**Fig. 4G**). We then used an *in vitro* culture assay to compare transcript expression of effector and exhaustion markers (*Pdcd1*, *Lag3*, *Havcr2*, *Ifng*) and memory-associated markers (*Tcf7*, *Il7r*) in B^+/+^ and B^-/-^ OT-I T cells under acute and chronic stimulation conditions. As expected, chronic stimulation led to increased *Pdcd1*, *Lag3*, and *Havcr2* expression and reduced *Tcf7* and *Il7r* expression in B^+/+^ OT-I cells compared to acute stimulation (**Fig. 4H**). However, under both stimulation conditions, B^-/-^ OT-I cells exhibited reduced expression of *Pdcd1*, *Lag3*, *Havcr2*, and *Ifng*, while *Tcf7* and *Il7r* expression remained significantly elevated relative to B^+/+^ OT-I cells. Collectively, these results indicate that Bhlhe40 is required for proper transcriptional programming and functional differentiation of effector and exhausted CD8 T cells.

### Bhlhe40 Sustains Glycolytic and Mitochondrial Metabolic Fitness in CD8 T Cells

Given that effector differentiation and cytokine production are tightly linked to metabolic rewiring, we next examined whether Bhlhe40 influences the metabolic profile of intratumoral CD8 T cells in 1956 tumor-bearing mice. Pathway enrichment analysis of genes correlated with *Bhlhe40* highlighted glycolysis-associated metabolic programs (**Figs. 4A**, **S7A**, and **S7B**), consistent with the metabolic rewiring of activated CD8 T cells. Key glycolytic genes such as *Aldoa*, *Gapdh*, *Pgk1*, and *Eno1,* were markedly reduced in anti-PD-1- or anti-CTLA-4- treated B^ΔCD8^ mice compared to B^f/f^ mice, particularly within the effector/exhausted clusters that also display the highest *Bhlhe40* expression (**Figs. 5A**, **S3**, and **S9A**). GSEA also showed negative enrichment of glycolysis-associated gene sets across several effector clusters in B^ΔCD8^ mice treated with anti-PD-1 or anti-CTLA-4 relative to B^f/f^ counterparts **(Figs. 5B**, **S5**, and **S6)**. Notably, in Bhlhe40^ΔT^ mice lacking Bhlhe40 in both CD4 and CD8 T cells, glycolytic gene expression was more severely impaired than in B^ΔCD8^ mice, suggesting a role for Bhlhe40 in CD4 T cells in facilitating CD8 T cell glycolytic fitness (**Fig. S9B**). Although we did not observe as extensive a reduction in OXPHOS-related transcripts in Bhlhe40-deficient CD8 T cells (**Fig. S9C**), GSEA revealed significantly decreased enrichment of OXPHOS-associated gene sets in several CD8 T cell clusters from anti-PD-1- and anti-CTLA-4- treated B^ΔCD8^ mice (**Figs. S5** and **S6**).

**Figure 5.**
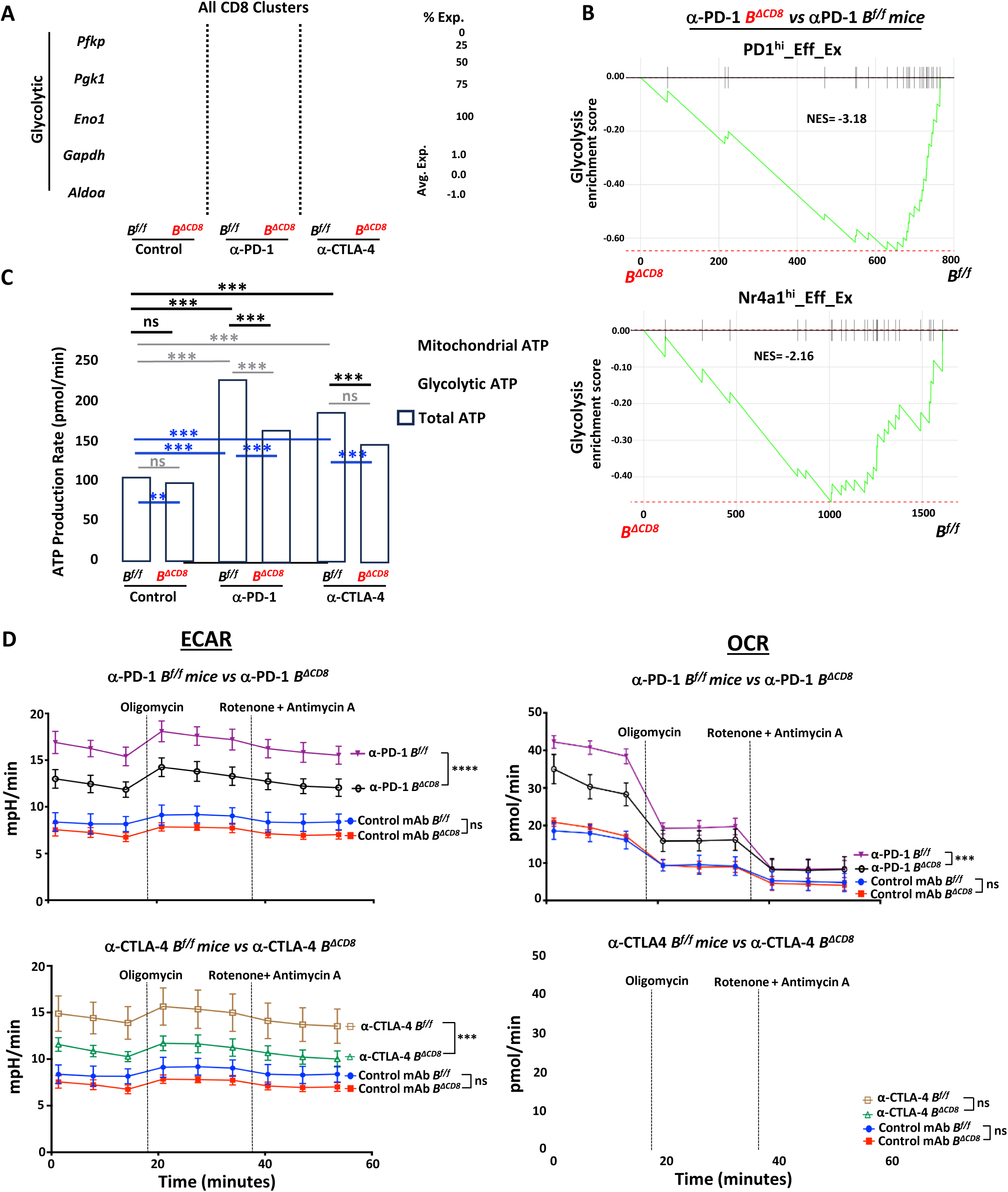
Bhlhe40 supports glycolysis and oxidative phosphorylation in CD8 T cells in a therapy-dependent manner. **(A)** Dot plot showing expression of select glycolytic genes (*Aldoa, Gapdh, Eno1,* etc.) in all CD8 T cell clusters. **(B)** GSEA showing enrichment of Hallmark glycolysis pathway in PD1^hi^_Eff_Ex and Nr4a1^hi^_Eff_Ex cluster between B^f/f^ vs B^ΔCD8^ mice with anti-PD-1 treatment. **(C)** XF real-time ATP production rate comparing glycolytic, mitochondrial, and total ATP on *ex vivo* intratumoral CD8 T cells in B^f/f^ and B^ΔCD8^ mice treated with control Ab, anti-PD-1, or anti-CTLA-4. **(D)** XF real-time ATP Seahorse assay showing ECAR and OCR on *ex vivo* intratumoral CD8 T cells in B^f/f^ and B^ΔCD8^ mice treated with control Ab, anti-PD-1, or anti-CTLA-4. For statistical comparisons in **(C)**, grey: mitochondrial ATP, blue: glycolytic ATP, and black: total ATP. **(C)** and **(D)** are representative of two independent experiments (∗∗p < 0.01; ∗∗∗p < 0.001; ns, not significant, unpaired t-test or 2-way ANOVA).

To determine whether these transcriptional changes corresponded to functional metabolic alterations, we performed Seahorse extracellular flux analysis on intratumoral CD8 T cells. Consistent with our scRNAseq-based indications, Bhlhe40-deficient CD8 T cells from anti-PD-1-treated B^ΔCD8^ mice exhibited significantly reduced ATP production from both glycolysis and mitochondrial OXPHOS (**Fig. 5C**) and ECAR/OCR time-course analyses similarly revealed diminished glycolytic activity and impaired OXPHOS in B^ΔCD8^ mice (**Fig. 5D**). In contrast, CD8 T cells from anti-CTLA-4-treated B^ΔCD8^ mice showed a significant reduction in glycolytic ATP production while largely maintaining mitochondrial ATP production (**Figs. 5C** and **5D**). These findings reveal that CD8 T cell-intrinsic Bhlhe40 is indispensable for maintaining glycolytic and mitochondrial ATP-generating metabolic pathways under anti-PD-1 therapy, whereas anti-CTLA-4 treatment uncovered a more selective reliance on Bhlhe40 for glycolysis but not mitochondrial respiration.

### T Cell-Intrinsic Bhlhe40 Sustains Anti-CTLA-4-Driven Th1 Responses and Tumor Rejection

Having defined the CD8 T cell defects caused by Bhlhe40 loss, we next asked whether differences in CD4 T cell responses could explain the divergent therapeutic outcomes, namely, that anti-CTLA-4 remains effective in B^ΔCD8^ mice but fails in B^ΔT^ mice. Notably, in both the 1956 and Y1.7LI tumor models, anti-CTLA-4 requires contributions from both CD4 and CD8 T cells for tumor control^30,41^. Therefore, the preserved efficacy of anti-CTLA-4 in B^ΔCD8^ mice cannot be interpreted as CD8 T cell dispensability; rather, it suggests that CD4 T cell-driven effects may compensate for Bhlhe40 loss in CD8 T cells. Consistent with prior studies^20,21,30,41^, anti-CTLA-4 more profoundly affected conventional CD4 T helper cells (CD4 T cells) and regulatory T cells (Tregs) relative to anti-PD-1 (**Figs. S10A** and **S10B**). Whereas anti-PD-1 did not alter the frequency of CD4 T cells in either B^f/f^ mice or B^ΔCD8^ mice, anti-CTLA-4 increased the frequency in both genotypes (**Fig. S10A**). As expected with anti-CTLA-4 clone 9D9 ^68^, treatment reduced Tregs, an effect that was independent of Bhlhe40 expression in CD8 T cells, as measured by both flow cytometry and scRNA-seq (Treg_1 cluster) **(Figs. 6A, S10A,** and **S10B)**. We next examined effector CD4 T cell populations that might contribute to anti-CTLA-4-mediated tumor control. scRNA-seq analysis identified several clusters of effector CD4 T cells, including multiple T helper 1 (Th1) populations characterized by *Ifng*, *Tbx21* (Tbet), and *Bhlhe40* expression **(Figs. 6B, S10C,** and **S10D)**. Amongst these, ICOS**^hi^**Bhlhe40**^hi^**_Th1_6 expressed the highest levels of *Bhlhe40, Tnf,* and *Ifng* among CD4 T cells, along with *Icos* (under anti-CTLA-4 treatment), *Tbx21*, and *Pdcd1* **(Figs. 6B, S10C,** and **S10D)**. This cluster also expressed *Bcl6*, indicating some transcriptional features of T follicular helper cells (Tfh) **(Fig. S10D)**. Gzmk^+^_Th1_8 expressed Th1 markers (e.g., *Tbx21*, *Ifng*) but also displayed high expression of *Gzmk*, along with moderate expression of *Tcf7* and *Slamf6* **(Fig. S10D)**. Comparative analysis with published datasets^45,69^ indicated that these murine Th1 subsets share transcriptional features with cytotoxic or highly activated effector populations in human cancer (**Fig. 6C**).

**Figure 6.**
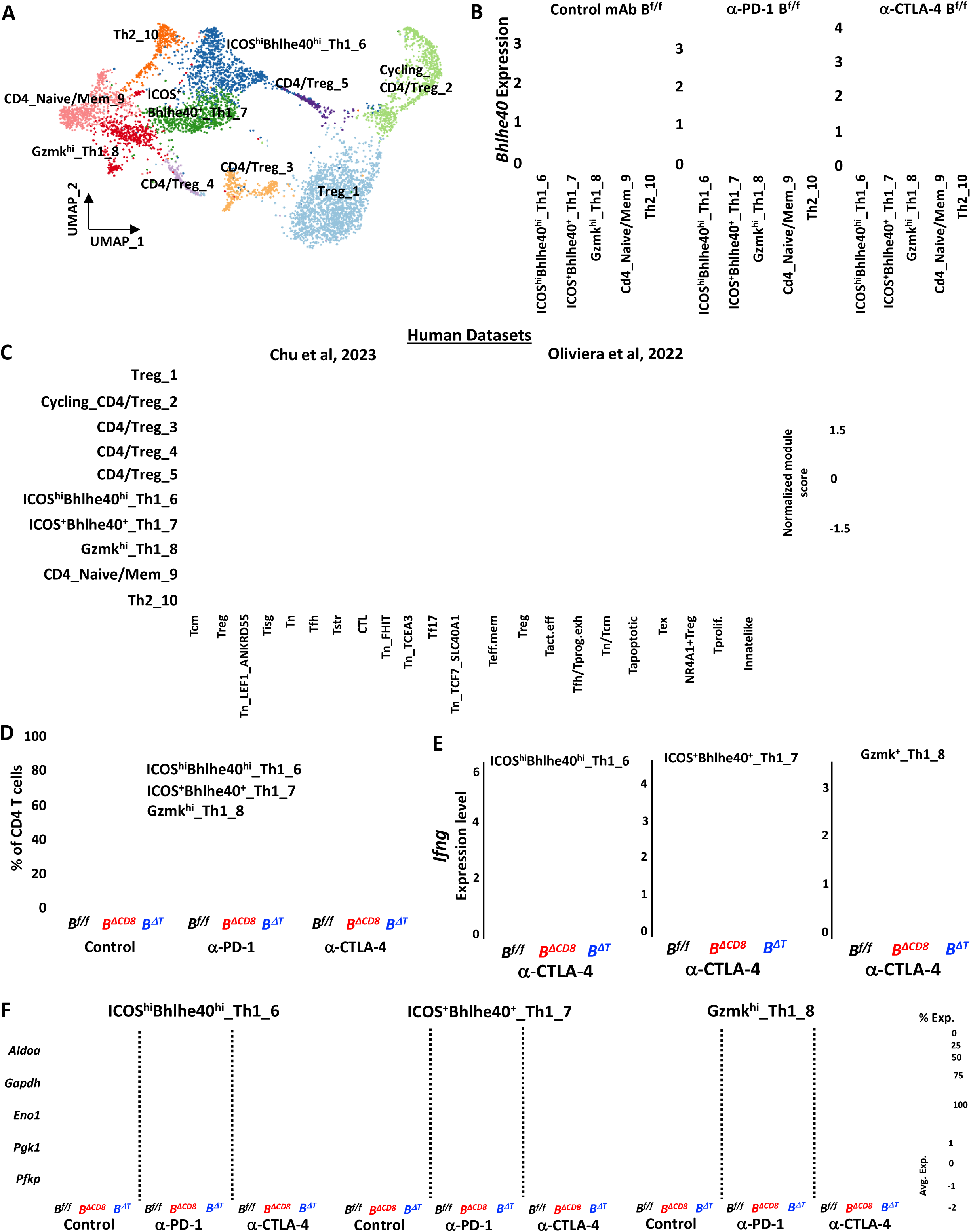
CD4 T cell-intrinsic Bhlhe40 sustains anti-CTLA-4-driven Th1 responses. **(A)** UMAP generated from integration of CD4 T cells depicting conventional CD4 T cell, Treg, and mixed conventional CD4 T cell and Treg clusters. **(B)** Violin plots showing *Bhlhe40* expression in CD4 T cells by cluster and treatment in B^f/f^ mice. **(C)** Heatmaps comparing features (module scores) of CD4 T cell clusters (rows) to published human CD4 T cell gene signatures (columns) identified/annotated by Chu et al. and Oliveira et al. **(D)** Stacked bar graph showing Th1 clusters as a percentage of CD4 T cells from B^f/f^, B^ΔT^ and B^ΔCD8^ mice treated with control Ab, anti-PD-1, or anti-CTLA-4. **(E)** Violin plot showing *Ifng* expression in ICOS^hi^Bhlhe40^hi^_Th1_6, ICOS^+^Bhlhe40^+^_Th1_7, and Gzmk^+^_Th1_8 CD4 clusters by treatment in B^f/f^, B^ΔT^, or B^ΔCD8^ mice. **(F)** Dot plot displaying select glycolytic transcript expression level and percent of cells in CD4 Th1 clusters.

As expected^20,41,70–72^, anti-CTLA-4 promoted expansion of ICOS**^hi^**Bhlhe40**^hi^**_Th1_6 and ICOS**^+^**Bhlhe40**^+^**_Th1_7 **(Figs. 6D** and **S10B)**. These populations expanded not only in B^f/f^ and B^ΔCD8^ mice, but also in B^ΔT^ mice, although the magnitude of expansion varied by genotype. Despite this expansion, several transcriptomic alterations were observed in mice with disrupted Bhlhe40 expression in both CD8 and CD4 T cells (B^ΔT^ mice) **(Figs. S10C** and **S10D)**. Notably, when comparing *Ifng* expression within CD4 T cells across treatments and knockout conditions, two important patterns emerged. First, anti-CTLA-4 induced a higher frequency of Th1 CD4 T cells expressing elevated levels of *Ifng* compared to anti-PD-1, and this increase was observed in both B^f/f^ and B^ΔCD8^ mice—treatment settings in which anti-CTLA-4-driven tumor rejection occurs (**Figs. 2B, 6D, 6E, S10C,** and **S10D**). Second, in the absence of Bhlhe40 in CD4 T cells (B^ΔT^ mice), *Ifng* expression was markedly reduced under all treatment conditions, including anti-CTLA-4, and in these mice, the efficacy of anti-CTLA-4 (as well as anti-PD-1) was lost (**Figs. 2B, 6D, 6E, S10C,** and **S10D**).

Interestingly, we observed a modestly decreased expression of glycolytic transcripts in CD4 effector T cells from B^ΔCD8^ mice (**Fig. 6F**). In contrast, CD4 T cells from B^ΔT^ mice, lacking Bhlhe40 in both CD4 and CD8 T cells, displayed striking reductions in glycolytic transcript expression compared to B^f/f^ mice, including in anti-CTLA-4-treated conditions. These findings suggest that CD8 T cell-intrinsic Bhlhe40 contributes indirectly to maintaining glycolytic transcript expression in CD4 effector T cells, whereas loss of Bhlhe40 across all T cell compartments leads to a more profound defects in metabolic transcript expression.

### T Cell-Intrinsic Bhlhe40 Shapes Macrophage Phenotypes

Our findings thus far indicate that Bhlhe40 orchestrates CD8 and CD4 T cell effector programs that are differentially required for anti-PD-1 and anti-CTLA-4 efficacy. Because we previously showed that anti-PD-1 and anti-CTLA-4 promote T cell-driven intratumoral macrophage remodeling, shifting from M2-like CX3CR1^+^ macrophages to M1-like iNOS^+^ macrophages^21,41,73^, we asked whether T cell-intrinsic Bhlhe40 also governs ICT-driven changes in intratumoral macrophages. We identified by scRNAseq multiple monocyte/macrophage clusters, including CX3CR1^+^ macrophage clusters expressing *Cx3cr1* and *Mrc1* (CD206) as well as distinct clusters expressing *Nos2* (iNOS) (**Figs. S11A-S11B**). Both scRNAseq and flow cytometry revealed a marked impairment in ICT-driven macrophage remodeling in B^ΔCD8^ mice, and an even more pronounced defect in B^ΔT^ mice lacking Bhlhe40 in both CD4 and CD8 T cells (**Figs. S11C-S11E**). Whereas anti-PD-1 or anti-CTLA-4 reduced the frequency of CX3CR1^+^ macrophages in B^f/f^ mice, this reduction was absent in B^ΔCD8^ and B^ΔT^ mice, which instead exhibited elevated frequencies of CX3CR1 macrophages under both control Ab and ICT conditions (**Figs. S11D** and **S11E**). This was also reflected by transcript expression of markers enriched on pro-tumoral macrophages (**Fig. S11C**), including *Trem2*, which we and others have shown is a therapeutic target on immunosuppressive macrophages^41,74,75^. Anti-PD-1 and especially anti-CTLA-4 expanded iNOS^+^ macrophages in B^f/f^ mice, whereas the frequency of iNOS^+^ macrophages was severely reduced in B^ΔCD8^ mice, with an even greater reduction occurring in B^ΔT^ mice (**Fig. S11E**). These findings indicate that Bhlhe40 expression in CD8 T cells, and more critically, in both CD4 and CD8 T cells, is required for ICT-induced macrophage remodeling, linking T cell-intrinsic Bhlhe40 to the shift from immunosuppressive CX3CR1 macrophages to proinflammatory iNOS macrophages within the TME.

### Bhlhe40 Correlates with Effector and Exhaustion Programs in Human Intratumoral CD8 T Cells

To relate our preclinical findings to human tumors, we next reanalyzed scRNAseq datasets^34,42,76,77^ from human cancer patients. These analyses revealed expression patterns between *BHLHE40* and several transcripts associated with T cell differentiation and functional programs similar to those observed in mice. In melanoma (Sade-Feldman et al. and Minowa et al.) and basal cell carcinoma (Yost et al.) datasets, gene-correlation analyses showed that *BHLHE40* expression was positively correlated with *TOX*, *PDCD1, ENTPD1, HAVCR2, LAG3, IFNG,* and *GZMB* (**Fig. 7A**). In contrast, *BHLHE40* exhibited negative correlation with *TCF7,* as well as *SELL* (CD62L) and *IL7R*. Consistent with these correlations, in triple-negative breast cancer (TNBC), *BHLHE40* was expressed in activated CD8 T cell clusters expressing *TOX* and *IFNG* (**Figs. S8C** and **S8D**).

**Figure 7.**
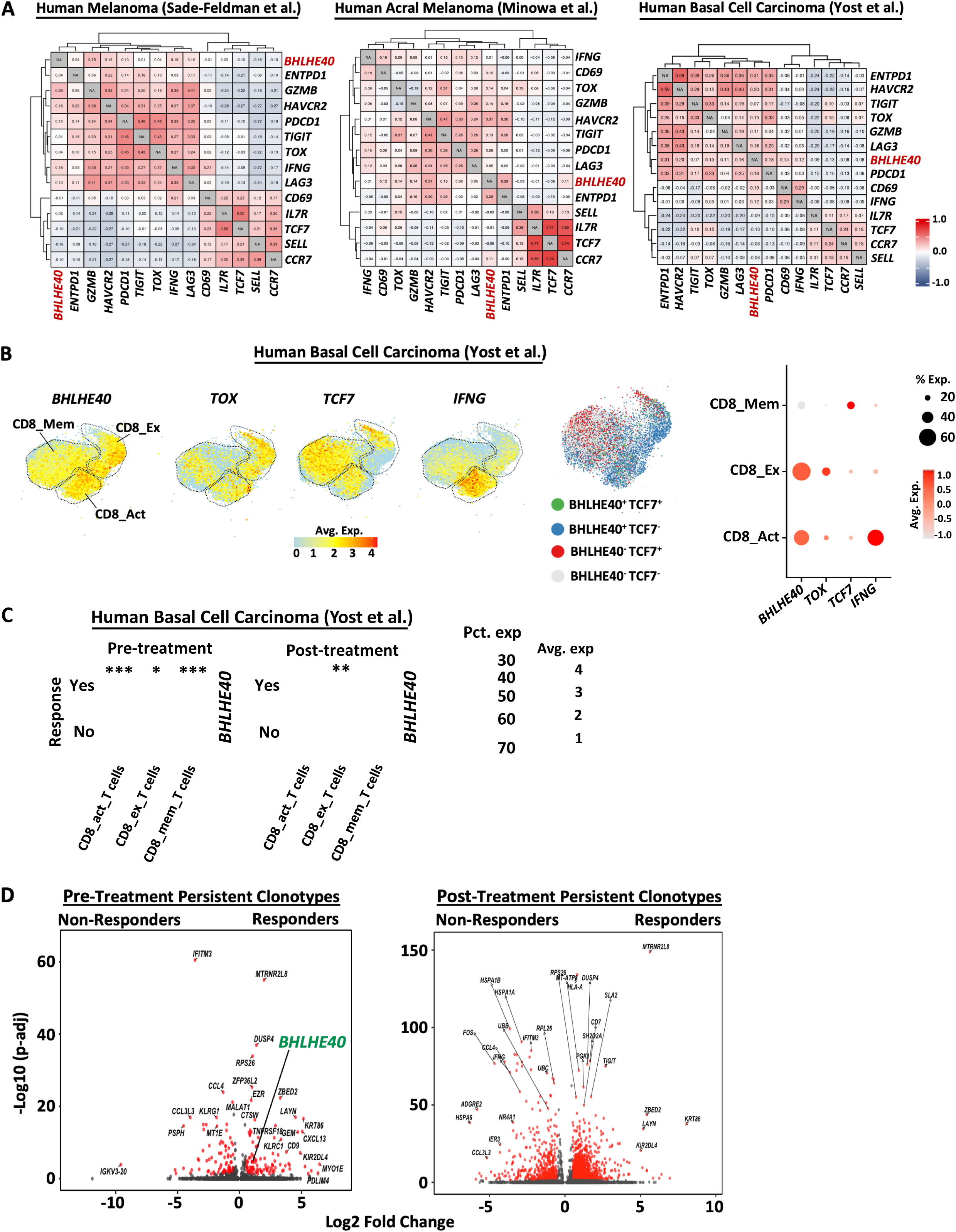
*BHLHE40* expression correlates with effector and exhaustion-associated programs in human intratumoral CD8 T cells. (**A**) Gene co-expression heatmap of human intratumoral CD8 T cells. (**B**) Feature plot showing expression of *BHLHE40, TOX, TCF7*, and *IFNG* in CD8 T cell clusters (left) from human basal cell carcinoma as identified and annotated by Yost et al., UMAP showing clusters positive or negative for *BHLHE40* and *TCF7* (middle), and dot plot displaying expression of select transcripts in CD8 T cell clusters (right). **(C)** Dot plot of CD8 T cells from pre- and post-treatment with anti-PD-1 basal cell carcinoma patients reveals elevated *BHLHE40* expression in activated and exhausted CD8 T cells. (**D**) Volcano plot showing differential expression of persistent clonotypes between responders and non-responders in pre- and post-treatment in Yost et al. Reanalysis of scRNAseq data from Sade-Feldman et al., Minowa et al., and Yost et al. shown in (**A**). Reanalysis of scRNAseq data from Yost et al shown in **(B-D)**. For **(C)**, *p < 0.05; ∗∗p < 0.01; ∗∗∗p < 0.001, Wilcoxon Rank-Sum test. For **(D)**, significantly differentially expressed transcripts (log_2_FC > 0.5, adjusted *P* value < 0.05) are shown in red. Statistical analysis was performed using a two-sided nonparametric Wilcoxon rank sum test with Bonferroni’s correction.

Next, we further analyzed the basal cell carcinoma dataset from Yost et al.^76^, which included pre-and post-treatment samples from responders and non-responders treated with PD-1 blockade. As expected, *BHLHE40* expression was highest in the CD8 T cell clusters classified by Yost et al. as ‘activated’ or ‘exhausted,’ whereas the ‘memory’ CD8 T cell cluster exhibited lower *BHLHE40* levels and higher *TCF7* expression (**Figs. 7B**). Notably, within these activated and exhausted CD8 T cell populations, *BHLHE40* expression was higher in responders compared with non-responders at pre-treatment (**Fig. 7C**). Following treatment, exhausted CD8 T cells showed elevated *BHLHE40* expression in responders. To determine whether *BHLHE40* expression associated with clinical outcome within stable TCR lineages, we analyzed persistent clonotypes present at both pre-treatment and post-treatment timepoints as defined in Yost et al. Persistent CD8 T cell clones from responders exhibited significantly higher *BHLHE40* expression compared to those from non-responders at pre-treatment (**Fig. 7D**). Collectively, our murine and human analyses identify Bhlhe40 as a central T cell-intrinsic regulator that couples effector and exhausted CD8 T cell differentiation and Th1 CD4 T cell function to macrophage remodeling and ICT efficacy, and further show that higher *BHLHE40* expression within tumor CD8 T cells associates with clinical response to PD-1 blockade. Further, these findings suggest that differential dependence on CD8 T cell-intrinsic Bhlhe40 contributes, at least in part, to the distinct mechanisms by which anti-PD-1 and anti-CTLA-4 mediate tumor rejection.

## DISCUSSION

Our study identifies Bhlhe40 as a transcriptional coordinator with distinct, therapy-specific requirements in CD4 and CD8 T cells during ICT. Mechanistically, Bhlhe40 promotes the acquisition of effector and exhausted CD8 T cell states and sustains IFN-γ production, while supporting both glycolytic and mitochondrial programs. Consistent with this, loss of Bhlhe40 skews CD8 T cells towards more Tpex-like states with elevated TCF-1 expression and reduced acquisition of terminal effector features, particularly under anti-PD-1 treatment and in the absence of ICT. Although the relationship between Bhlhe40 and several transcripts/proteins associated with CD8 T cell function may be indirect, some of the observed changes are likely attributable to Bhlhe40-mediated transcriptional activation or repression of specific target genes. For example, Bhlhe40 has been shown to positively regulate *Ifng* and negatively regulate *Il10*^23,24,28^, consistent with our findings. In addition, *Tcf7* contains several E-box motifs within its regulatory regions^78^ and CUT&RUN analyses of intratumoral CD8 T cells revealed Bhlhe40 enrichment at regulatory regions of *Tcf7* under control and anti-PD-1 treatment conditions. This raises the possibility that Bhlhe40 may not only be co-expressed with markers of activated, effector, or exhausted CD8 T cells due to shared upstream signals such as TCR stimulation, but may also contribute directly to driving CD8 T cell differentiation toward effector and exhausted phenotypes. Collectively, our findings support a model in which Bhlhe40 functions as a transcriptional switch that drives effector (and exhausted) phenotypes in tumors in part by modulating TCF-1 expression. Consistent with this model, Wu et al.^79^ recently demonstrated using in vitro chronic stimulation and chronic infection models that Bhlhe40 regulates a differentiation checkpoint between progenitor and intermediate exhausted CD8 T cells.

Anti-PD-1 and anti-CTLA-4 each utilize both shared and distinct mechanisms^20,21,41^. Anti-PD-1 predominantly expands and sustains effector function from a PD-1^+^ TCF-1 progenitor (Tpex) CD8 T cell pool, as terminally exhausted cells are epigenetically constrained and minimally reinvigorated^47^, whereas anti-CTLA-4, directly and via CD4 T cell help, enhances CD8 T cell priming/expansion with more memory-like and less terminally exhausted features (increased TCF-1 and decreased TOX)^11,20,21,58,60,80,81^.

These differences provide a basis for the differential requirement for Bhlhe40 in CD8 T cells during anti-PD-1 versus anti-CTLA-4 therapy. Whereas anti-PD-1 efficacy likely depends on Bhlhe40 to drive the transition of CD8 Tpex cells into effector (and exhausted) states, anti-CTLA-4 may rely less on CD8 T cell-intrinsic Bhlhe40 and instead promote expansion of CD8 T cells with more memory-like or less terminal effector/exhausted features. Nevertheless, loss of CD8 T cell-intrinsic Bhlhe40 did cause functional alterations, including reduced CD8 T cell-derived IFN-γ, during anti-CTLA-4 treatment, although anti-CTLA-4 still expanded Psmd6-specific CD8 T cells and maintained tumor rejection. This preservation of tumor control may be due in part to compensatory actions of CD4 T cells. We recently found that IFN-γ^+^ Th1-like ICOS^+^ CD4 T cells induced by anti-CTLA-4^20,71^ highly express Bhlhe40 ^41^, consistent with prior reports that Bhlhe40 promotes conventional Th1 differentiation and sustains IFN-γ production in autoimmune and infection settings^23,32,82^. These anti-CTLA-4-induced ICOS^+^ Th1-like cells likely contribute to tumor control in B^ΔCD8^ mice, where CD8 T cell function is compromised. In contrast, mice lacking Bhlhe40 in both CD4 and CD8 T cells failed to respond to either ICT modality, highlighting the cooperative roles of these subsets in orchestrating anti-tumor immunity. Together, these findings indicate that the contribution of Bhlhe40 to ICT responses is not uniform across T cell subsets or therapeutic modalities, but instead reflects differential mechanisms engaged by anti-PD-1 versus anti-CTLA-4.

Importantly, the phenotypic and functional impairments observed in CD8 T cells lacking Bhlhe40 were accompanied by defects in ICT-driven remodeling of the tumor myeloid compartment from CX3CR1^+^ macrophages to iNOS^+^ macrophages. These CX3CR1 macrophages likely constitute a large fraction of M2-like tumor-associated macrophages, often co-expressing CD206, C1q, ApoE, TREM2, etc., although marker usage and nomenclature varies across studies^14,19–21^, whereas iNOS^+^ macrophages are consistent with inflammatory M1-like macrophages. Notably, the relatively greater expansion of iNOS^+^ macrophages observed with anti-CTLA-4 compared to anti-PD-1 in CD8 T cell Bhlhe40-deficient mice may reflect CD4 T cell-derived IFN-γ and Th1-like help, whereas combined loss of Bhlhe40 in both CD4 and CD8 T cells profoundly impaired macrophage remodeling, even under anti-CTLA-4 therapy.

Our data also link Bhlhe40 to metabolic programming in a partially therapy-dependent manner. Under anti-PD-1 treatment, Bhlhe40-deficient CD8 T cells exhibited marked reductions in both glycolytic and OXPHOS capacity, with reduced mitochondrial respiration consistent with prior work implicating Bhlhe40 in sustaining mitochondrial fitness during anti-PD-1 therapy^26^. Under anti-CTLA-4 treatment, Bhlhe40-deficient CD8 T cells retained OXPHOS but displayed reduced glycolytic activity. Interestingly, loss of Bhlhe40 in CD8 T cells also impaired glycolytic transcript expression in CD4 effector T cells, reflecting alterations in CD8 T cells that either directly influence CD4 T cells or reshape the TME in ways that secondarily impact CD4 T cells. Notably, in B^ΔT^ mice lacking Bhlhe40 in both CD4 and CD8 T cells, glycolytic transcript expression in CD4 T cells was markedly reduced across treatment conditions. ICT-induced, Bhlhe40-dependent T cell-driven changes to tumor metabolism may also influence T cell metabolic fitness, as increased tumor glucose consumption and lactate production could further constrain T cell glycolysis by acidifying the TME^83^.

These observations align with the well-established requirement for robust glycolytic and mitochondrial programs to support effector cytokine production, cytotoxicity, and persistence of T cells in the TME^84–86^. In particular, glycolysis supports effector responses, whereas mitochondrial metabolism supports sustained function and long-term survival under conditions of chronic stimulation^83,85–87^. Given the requirement for T cells to upregulate glycolysis upon effector differentiation, it is notable that signals downstream of TCR activation and costimulation that drive metabolic reprogramming also induce Bhlhe40 expression, placing Bhlhe40 at the interface between activation-induced metabolic enhancement and effector programming.

Analysis of human cancer datasets revealed that *BHLHE40* is enriched in tumor-reactive and activated/exhausted CD8 T cells across multiple tumor types, inversely correlated with *TCF7*, and positively associated with *TOX, GZMB,* and *IFNG*. Moreover, analysis of the Yost et al. dataset^76^ revealed that persistent TCR clonotypes—defined as clones present at both pre- and post-treatment timepoints—exhibited higher *BHLHE40* expression in responders compared to non-responders before PD-1 blockade. These findings suggest that *BHLHE40* marks an effector-biased intratumoral CD8 T cell population with enhanced capacity to persist through therapy and sustain anti-tumor immunity. Together, these observations support the relevance of our murine findings to human ICT responses and suggests that Bhlhe40 expression could serve as a biomarker for T cell differentiation state and functional competence in the TME.

Collectively, our findings position Bhlhe40 as a subset- and therapy-specific regulator of ICT responses, integrating effector programming, metabolic fitness, and myeloid remodeling to provide mechanistic insight into the distinct modes of action of anti-PD-1 and anti-CTLA-4. While multiple transcription factors influence T cell programs relevant to ICT, our study provides one of the clearest demonstrations of a transcriptional regulator that imposes T cell subset-specific requirements for anti-PD-1 versus anti-CTLA-4 efficacy, resulting in complete loss of one therapy while fully sparing the other. From a translational perspective, these results suggest that strategies to sustain or enhance Bhlhe40 activity, particularly in CD8 T cells during PD-1 blockade, may improve therapeutic efficacy. Conversely, the ability of CD4 T cells to preserve anti-CTLA-4 efficacy despite CD8 T cell impairment suggests opportunities to harness or amplify these complementary pathways to improve immunotherapy responses.

## MATERIALS AND METHODS

### Mice

All mice used were on a C57BL/6 background. Wild type C57BL/6J, *CD4-Cre* (B6.Cg-Tg(Cd4-cre)1Cwi/BfluJ)^36^: JAX stock #022071, *CD8 E8I-Cre* (B6.Tg(Cd8a-cre)1Itan/J)^37^: JAX stock #008766, *Bhlhe40^-/-^* (B6.129S1(Cg)-Bhlhe40tm1.1Rhli/MpmJ): JAX stock #029732, and OT-I TCR transgenic (Tg(TcraTcrb)1100Mjb/J): JAX stock #003831 mice were purchased from Jackson Labs. All *in vivo* experiments used 8–12-week-old female or male (to match the sex of the tumors), all mice were housed in our specific pathogen-free animal facility. *Bhlhe40^f/f^*mice have been previously reported^23,24,30,88^. For experiments using conditional Bhlhe40 KO mice, *Bhlhe40^f/f^* littermates or *CD8 E8I-Cre Bhlhe40^+/+^* mice were used as controls. All studies were performed in accordance with procedures approved by the Institutional Animal Care and Use Committee (IACUC) at The University of Texas MD Anderson Cancer Center, an AAALAC accredited institution.

### Tumor lines

The 1956 MCA-induced sarcoma line used in this study was generated in female wild type C57BL/6 mice and were banked as low-passage tumor cells as previously described^89^ and were obtained from Robert Schreiber (Washington University in St. Louis School of Medicine). Y1.7LI is a melanoma line derived from the Braf^V600E^ Cdkn2a^-/-^ Pten^-/-^ YUMM1.7 line^40^ engineered to express mLama4, an H2-K^b^-restricted MHC class I neoantigen, and mItgb1, an I-A^b^-restricted MHC class II neoantigen, as previously described^41^. Tumor lines from the same cryopreserved stocks that were used in this study tested were authenticated and found to be free of non-mouse cells by STR profiling (IDEXX CellCheck^TM^ mouse 19 plus *Mycoplasma* spp. testing) and confirmed free of common mouse pathogens and mycoplasma by IDEXX IMPACT^TM^ I mouse pathogen testing.

### Tumor transplantation

1956 MCA sarcoma cells, B16-F10-OVA (B16-OVA) melanoma cells, and Y1.7LI melanoma cells were from frozen stocks and propagated in R-10 plus BME media [RPMI media (Hyclone) supplemented with 1% L-glutamine, 1% penicillin-streptomycin, 1% Na Pyruvate, 0.5% Na Bicarbonate, 0.1% 2-Mercaptoethanol, and 10% FCS (Hyclone)]. Upon thawing, tumor lines were passaged 3-5 times before experimental use. Prior to injection, cells were washed extensively, resuspended at a concentration of 6.67 × 10^6^ cells per mL in endotoxin-free PBS for 1956 and at a concentration of 3.33 × 10^6^ cells per mL in endotoxin-free PBS for B16-OVA and Y1.7LI and then 150 μl (1 × 10^6^ cells per mouse for 1956 and 0.5 × 10^6^ cells per mouse for B16-OVA and Y1.7LI) injected subcutaneously into the flanks of recipient mice. Tumor cells were >90% viable at the time of injection as assessed by trypan blue exclusion. Tumor growth was quantified by caliper measurements and expressed as the average of two perpendicular diameters.

### Tumor harvest

Established tumors were isolated from mice, minced and treated with 1 mg/ml type IA collagenase (Sigma) in HBSS (Hyclone) for 45 minutes at 37 °C. Cells were washed thrice. Red blood cells were lysed using ACK lysis buffer (Gibco). To remove aggregates and clumps, cells were passed through a 40-micron strainer.

### Immune checkpoint therapy

Mice were treated intraperitoneally with 200 μg of anti-PD-1 or anti-CTLA-4 on days 3, 6, 9 post-tumor transplant for 1956 and on days 7, 10, 13, 16, and 21 for Y1.7LI. For controls, mice were injected with 200 μg of isotype control antibodies.

### *In Vivo* antibodies

Anti-PD-1 (rat IgG2a clone RMP1-14) and anti-CTLA-4 (murine IgG2b clone 9D9) immune checkpoint antibodies were purchased from Leinco Technologies. Isotype control antibodies (rat IgG2a clone 1-1, mouse IgG2a clone OKT3, and rat IgG2b clone 1-2) were purchased from Leinco Technologies. All *in vivo* antibodies purchased from Leinco were “*In vivo* Platinum”-grade and were verified free of mouse pathogens (IDEXX IMPACT^TM^ I mouse pathogen testing).

### Quantitative RT-PCR

RNA was extracted from sorted T cells using RNAeasy Plus Mini Kit (Qiagen). 1 μg of RNA was reverse-transcribed and subjected to qPCR using the SuperScript III Platinum Two-Step qRT-PCR Kit with SYBR Green (Invitrogen). qPCR was performed on the StepOne Real-Time PCR System (Applied Biosystems). Each sample was run in triplicate for each gene and the cDNA from each sample was divided equally per reaction in a 20 μl volume. The qPCR conditions were as follows: 50°C for 2 minutes and 95°C for 2 minutes, followed by 40 cycles of 95°C for 15 seconds and 60°C for 30 seconds. Melting curve analysis was performed to confirm a single amplicon. Differences in gene expression were determined using the equation 2^-ΔΔCt^, where the Ct value for *Gapdh* control was subtracted from the Ct value of *Bhlhe40* the to yield the ΔCt value. For each sample, the ΔCt value *Bhlhe40* done in triplicate was averaged and compared to give one ΔΔCt value per sample. Mouse quantitative RT-PCR primers for *Bhlhe40* were as follows: forward primer-5’ ACGGAGACCTGTCAGGGATG3’ and reverse primer-5’GGCAGTTTGTAAGTTTCCTTGC3’. Mouse quantitative RT-PCR primers for *Gapdh* were as follows: forward primer-5’AGGTCGGTGTGAACGGATTTG3’ and reverse primer-5’TGTAGACCATGTAGTTGAGGTCA3’.

### Single-cell RNA Sequencing

#### Antibody hashing for multiplexing and single-cell RNA sequencing library preparation

Antibody hashing and multiplexing was utilized for scRNAseq. For the scRNAseq Experiment #1 and Experiment #2, cell labeling was performed according to an adapted BioLegend cell hashing protocol (TotalSeq™-C Antibodies and Cell Hashing with 10x Single Cell 5’ Reagent Kit v1.1 Protocol, BioLegend). Single cell suspensions of harvested tumors from treated mice of different groups were resuspended in BioLegend Cell Staining Buffer containing Fc block and then surface stained with an antibody master mix consisting of anti-CD45.2/PE, anti-CD90.2/Thy1.2-PE-Cy7, and live/dead dye (NIR) and then barcoded antibodies with unique hashtags were added according to Supplement Table 1. Hashtag antibodies were used at a concentration of 1 μg per 2 million cells. Cells were then washed 3X with BioLegend Cell Staining Buffer and pooled for single-cell library generation and CITE-seq (cellular indexing of transcriptomes and epitopes by sequencing) through multiplexing. Cells were counted on a Countess 3 FL automated cell counter (Life Technologies) and viabilities were determined using trypan blue exclusion assay. Cell capture processing and gene expression and feature barcode library preparations were performed following 10X Genomics’ guidelines for 5’ scRNAseq by encapsulating sorted live CD45^+^ tumor infiltrating cells [CG000330_Chromium Next GEM Single Cell 5’ v2 (Dual Index) with Feature Barcode technology-Rev F]. QC steps after cDNA amplification and library preparation steps were carried out by running ThermoFisher Qubit HS dsDNA Assay along with Agilent (Santa Clara, CA) HS DNA Bioanalyzer for concentration and quality assessments, respectively. Library sample concentrations were verified using qPCR using a KAPA Biosystems KAPA Library Quantification Kit prior to pooling. Libraries were normalized to 5 nM for pooling. The pool was sequenced using a NovaSeq6000 S4-XP, 200-cycle flow cell lane. The run parameters used were 26 cycles for read 1, 90 cycles for read2, 10 cycles for index1, and 10 cycles for index2 as stipulated in the protocol mentioned above.

#### Single-cell RNA sequencing alignment, barcode assignment, and UMI Counting

The Cell Ranger Single-Cell Software Suite available at https://support.10xgenomics.com/single-cell-gene-expression/software/overview/welcome was used to perform sample demultiplexing, barcode processing and single-cell 5’ counting. “Cellranger mkfastq” was used to demultiplex raw base call files from the NovaSeq6000 sequencer, into sample-specific fastq files. Files were demultiplexed with 81.9% to 97.1% perfect barcode match, and 90%+ q30 reads. Afterward, fastq files for each sample were processed with “Cellranger count”, which was used to align samples to mm10 genome, filter and quantify. For each sample the recovered-cells parameter was specified as 10,000 cells that we expected to recover for each individual library.

#### Preprocessing analysis with Seurat package

The Seurat pipeline was applied to each dataset following tutorial specifications from [https://satijalab.org/seurat/articles/archive; version 3.2] and [https://hbctraining.github.io/scRNA-seq_online/]. QC process was performed for each experiment and group separately. Genes that were expressed in less than 3 cells and cells that contained less than 500 transcripts (UMI’s), or had less than 250 and more than 6500 genes, or more than 10% of mitochondrial transcripts, were excluded. Data were normalized using LogNormalize method. Potential doublets were removed by using scDblFinder package (1.16.0). For generating integrated CD8 T cell object, we used gene expression: *Cd3e > 0*, *Cd8a* > 0 & *Cd4* = 0, similar for generating CD4 T cell object we used: *Cd3e > 0, Cd4* > 0 & *Cd8a* = 0. All Cd8 and Cd4 data from all groups and experiments were integrated into a single CD8 or CD4 Seurat object. Expression matrix from all group were integrated using batch effect correction algorithm implemented in the Seurat V5 pipeline, employing anchor-based reciprocal principal component analysis (RPCA). Briefly, repeatedly variable genes across groups were selected by using ‘SelectIntegrationFeatures’ function. ScaleData and RunPCA were performed using these genes. Next, anchors were identified with ‘FindIntegrationAnchors’ and integration was performed using these anchors through ‘IntegrateData’. Unsupervised graph-based clustering was then performed on the integrated dataset. A UMAP dimensional reduction was performed on the scaled matrix (with most variable genes only) using first 40 PCA for CD8 and 50 PCA for CD4 components to obtain a two-dimensional representation of the cell states. For clustering, we used the function FindClusters that implements SNN (shared nearest neighbor) modularity optimization-based clustering algorithm.

#### Identification of cluster-specific genes and marker-based classification

To identify marker genes, the FindAllMarkers function was used with likelihood-ratio test for single cell gene expression. For heatmap representation, average expression of markers inside each cluster was used. To compare gene expression for the clusters inside cohorts (e.g., T cells, macrophages) we used FindMarkers function to calculate average log2 fold change and identify differentially expressed genes between each pair of experimental conditions using a Wilcoxon Rank-Sum test for calculating p-values and Bonferroni correction for adjusted p-values. Gene set enrichment analysis was performed using the fgsea package (version 1.32.0) to assess the enrichment of Hallmark gene sets from the Molecular Signatures Database (MSigDB). Hallmark gene sets were obtained using the msigdbr package (version 7.5.1). Differential gene expression analysis was performed using the FindMarkers function in Seurat (version 5.1.0) with the following parameters: threshold of log2 fold change = 0.2, and minimum percentage of expression = 0.1. Genes were ranked according to their log2 fold change, and these rankings were used as input for the enrichment analysis using the fgsea package.

#### Comparison to published datasets

The gene lists defining phenotype of intratumoral T cells were retrieved from published human or mouse scRNAseq datasets^34,42,44,45,69,90^ (**Tables S2-S5**) to compare the cluster annotation between published study and our current study. The module scores for individual cells were calculated using the “AddModuleScore” function in the Seurat package. The results were visualized using the normalized average module scores for each cluster (**Figs. 2H**, and **6C**).

#### Analysis of Bhlhe40 Regulon Activity

Single-Cell Regulatory Network Inference and Clustering (SCENIC) (version 1.3.1)^91^ was performed with default parameters to explore transcriptional factor (TF) regulatory networks. Murine mm10 TF annotations were used as a reference, and co-expression modules between TFs and candidate target genes were inferred using GENIE3. Using the RcisTarget package (version 1.20.0), TF binding motifs enriched in the target gene sets were identified. Cell-type specific regulators were identified based on the Regulon Specificity Score (RSS)^92^.

#### Trajectory analysis

Trajectory analysis was performed using the Monocle3 package (version 1.3.7)^93^. The Seurat object was converted into the Monocle3 object using the “as.cell_data_set” function. Cluster identities, dimensional reduction, and UMAP coordinates were transferred from Seurat object into the Monocle3 object. The trajectory was inferred with the default parameters.

### Analysis of human datasets

The human scRNAseq data were collected from previously published datasets. Read count matrix was downloaded and metadata generated in Yost et al manuscript^76^ (GSE123813). For Bassez et al^77^ read count matrix and metadata are downloaded through VIB-Ku LEUVEN center for cancer biology (https://lambrechtslab.sites.vib.be/en/single-cell). Read count matrix from Sade-Feldman et al.^42^ dataset was downloaded through GSE120575. Seurat objects then were generated and processed with Seurat pipeline. Seurat object of CD8 T cells in the paper from Minowa et al.^90^ (GSE242477) were download from Zenodo (https://doi.org/10.5281/zenodo.12761836). For volcano plot from Oliveira et al^34^, DEG list provided in the Oliveira et al. paper were used (phs001451.v3.p1). For volcano plot from Minowa et al, DEGs were identified by using FindMarkers function. For cell-type specific co-expression, inference from scRNAseq data, CS-CORE package was utilized to create the co-expression heatmap^94^.

### Ex Vivo T Cell Stimulation

For PMA/ionomycin T cell stimulation, cells from tumors were enriched for CD8 T cells using Miltenyi mouse CD8^+^ T cell enrichment from TIL kit by following manufacturer’s protocol. Cells were then incubated with 10 ng/ml of PMA and 1 μg/ml of ionomycin in the presence of BD GolgiPlug (BD Bioscience) for 6 hours at 37 °C. and stained for IFN-γ and TNFα.

### Adoptive Transfer and OT-I CD8 T cell Acute and Chronic Stimulation Assay

For adoptive transfer of OT-I CD8 T cells (transgenic for a TCR recognizing the H2-K^b^-presented SIINFEKL epitope of OVA) into mice bearing B16-OVA melanoma, splenic CD8 T cells from B^+/+^ OT-I and B^-/-^ OT-I mice were stimulated with 1 nM of SIINFEKL peptide and intravenously transferred (4 × 10^6^ cells per mouse). For acute and chronic antigen stimulation, naïve OT-I CD8 T cells from B^+/+^ OT-I and B^-/-^ OT-I mice were cocultured with naïve splenocytes pulsed with 1 nM of SIINFELK peptide. Chronic antigen stimulation was done to OT-I CD8 T cells by stimulating cells every 2 days with SIINFEKL peptide (1 nM) and rIL-2. Cells were harvested on day 10 post-first stimulation and processed for RT-qPCR.

### Tetramer staining

Mutant Psmd6-H2-K^b^-PE tetramers (IAPYYEAL) were obtained from the Baylor College of Medicine MHC Tetramer Production Facility. Cells were incubated at room temperature for 5 minutes with 500ng of rat anti-mouse anti-CD16/32 (mouse Fc block; clone 2.4G2, BD Biosciences) at 1µg/million cells. Peptide-H2-K^b^ tetramers conjugated with PE for mutated Psmd6 (1:500 dilution) were added to the cells and incubated for 20 minutes at 37°C. Cells were further stained with an antibody mixture cocktail: anti-mouse CD45 BV605 (clone 30-F11) BioLegend (1:800 dilution), anti-mouse Thy1.2-PE-Cy7 (clone 30-H12) BioLegend (1:500 dilution), anti-mouse CD4-BV711 (clone RM4-5) (1:200 dilution) BioLegend, anti-mouse CD8a-BV786 (clone 53-6.7) BioLegend (1:200 dilution), and live/dead (Zombie NIR) in 100 μl FACS buffer for 20 minutes at 4°C.

### Flow cytometry

For flow cytometry, cells were stained for 5 minutes at room temperature with Fc block (anti-CD16/32) clone 2.4G2 (BD Bioscience) at 1 μg/million cells and then surfaced stained for 20 minutes at 4 °C. Surface antibodies were diluted in FACS staining buffer (PBS with 2% FCS, 2 mM EDTA, and 0.05% NaN_3_ (Sigma)). Anti-mouse CD45 BV605 (clone 30-F11) BioLegend (1:800 dilution), anti-mouse CD3e-APC (clone 145-2C11) BioLegend (1:200 dilution), anti-mouse CD4-BV711 (clone RM4-5) (1:200 dilution) BioLegend, anti-mouse CD8a-BV786 (clone 53-6.7) BioLegend (1:200 dilution), anti-mouse CX3CR1-FITC (clone SA011F11) BioLegend (1:1000), anti-mouse I-A/I-E-BV650 (clone M5/114.15.2) BioLegend (1:3000 dilution), anti-mouse CD24-BV711 (clone M1/69) BD Bioscience, anti-mouse CD11c-BV786 (clone HL3) BD Bioscience (1:400 dilution), anti-mouse F4/80-BUV395 (clone T45-2342) BD Bioscience (1:400 dilution), anti-mouse CD64-BV421 (clone X54-5/7.1) BioLegend (1:400 dilution), anti-mouse Ly6G-Alexa Fluor 700 (clone 1A8) BD Bioscience (1:400 dilution), and anti-mouse CD11b-APC (clone M1/70) BioLegend (1:400 dilution) were used for surface stain at indicated dilutions. Zombie NIR Viability dye (BioLegend) was added at 1:500 during surface stain. For intracellular staining, surface-stained cells were fixed and permeabilized with BD fixation and permeabilization kit (BD Biosciences). Fixed and permeabilized cells were then stained with anti-mouse Mrc1 (CD206)-APC-Cy7 (clone C068C2) BioLegend (1:400 dilution) and anti-mouse iNOS/Nos2-PE (clone CXNFT) Thermo Fisher (1:400 dilution) for 30 minutes at 4 °C. For intracellular cytokine staining of lymphocytes, cells were isolated and then incubated at 37 °C for 6 hours with GolgiStop (BD Bioscience). Cells were then harvested, washed, stained for 5 minutes at room temperature with Fc block (anti-CD16/32) clone 2.4G2 (BD Bioscience) at 1 μg/million cells and then surfaced stained for 20 minutes at 4°C, and then fixed and permeabilized with BD fixation and permeabilization kit (BD Biosciences). Fixed and permeabilized cells were then stained with anti-IFN-γ-APC (XMG1.2) BioLegend (1:200 dilution) and anti-TNF-α-PE-Cy7 (clone MP6-XT22) BioLegend (1:200 dilution) for 30 minutes at 4°C. All flow cytometry was performed on the LSR Fortessa X-20 (BD Biosciences) and analyzed using FlowJo software (TreeStar).

### Fluorescence-activated cell sorting

For sorting of CD8/CD4 T cells and CD45.2^+^Thy1.2^+^ cells, intratumoral cells were stained for 5 minutes at room temperature with Fc block (anti-CD16/32) clone 2.4G2 (BD Bioscience) at 1 μg/million cells and then were stained with surface antibodies against CD45, CD90.2/Thy1.2, CD8a, and CD4 for CD4/CD8 sorting and with CD45.2-PE (clone 104) BioLegend (1:1000 dilution) and CD90.2/Thy1.2 for CD45.2^+^Thy1.2^+^ sorting along with Zombie NIR Viability dye and incubated for 30 minutes at 4°C. Cells were washed twice with MACS buffer and sorted gating on live CD45^+^, CD90.2/Thy1.2^+^, CD8a^+^, CD4^+^ or CD45.2^+^,Thy1.2^+^. Sorting was performed on a BD FACSAria II (BD Biosciences). CD8/CD4 sorted cells were processed for qPCR or CUT&RUN experiment and CD45.2^+^Thy1.2^+^ sorted cells were processed for scRNA sequencing. The purity of the sorted cells for scRNAseq was greater than 97% as assessed during post-sort cellular analysis.

### Magnetic cell sorting

Miltenyi Biotec’s magnetic cell separation was used to isolate intratumoral CD8 T cells for Seahorse real-time ATP rate assay. Single cell suspension from tumors were resuspended in MACS buffer, along with 10 μl of CD8 (TIL) microbeads per 10^7^ total cells and incubated for 15 minutes in 4°C. After incubation, CD8 T cell fractions were isolated in a magnetic field using MACS separator according to the manufacturer’s protocol. Isolated CD8 T cells were processed for Seahorse real-time ATP rate assay.

### Seahorse XF Real-Time ATP Rate Assay

The Seahorse XF Real-Time ATP Rate Assay (Agilent, 103592-100) were performed on a Seahorse XFe96 Analyzer (Agilent) according to the manufacturer’s protocol. Briefly, 1 × 10 cells/well were seeded in Seahorse XF RPMI-1640 medium (Agilent, 103576-100) supplemented with 1 mM pyruvate (103578-100), 10 mM glucose (103577-100), and 2 mM glutamine (103579-100) in a Cell-Tak–coated Seahorse XF96 cell culture microplate. Cells were centrifuged at 200 × g for 1 minute (no brake) and then incubated in a 37°C non-CO incubator for 30 minutes to allow complete adherence. Pre-hydrated sensor cartridges were loaded with 1.5 µM oligomycin and 0.5 µM rotenone/antimycin, which were sequentially injected during the assay to measure oxygen consumption rate (OCR) and extracellular acidification rate (ECAR) in real time. Data were analyzed using Wave Software (Agilent, v2.6.3.5) to calculate ATP production rates derived from mitochondrial oxidative phosphorylation (OXPHOS) and glycolysis, and values were normalized to seeded cell numbers.

### Cleavage Under Targets and Release Using Nuclease (CUT&RUN) Assay

CUT&RUN was performed at the MD Anderson Cancer Center Epigenomics Profiling Core following the protocol as described^95,96^ with some modifications. Briefly, nuclei isolated from ∼500,000 live intratumoral CD8 T cells were immobilized on Concanavalin A-coated magnetic beads (Bangs Laboratories) and permeabilized using wash buffer containing 0.02% digitonin (Promega). Bead-bound nuclei were incubated with rabbit IgG (Millipore) and Bhlhe40 (NB100-1800 EpiCypher) antibodies overnight at 4°C. Next day, targeted chromatin digestion was achieved by pAG-MNase (EpiCypher) binding for 10min at room temperature followed by incubation with CaCl_2_ at 4°C. CUT&RUN DNA was purified using MinElute columns (Qiagen). Library preparation was carried out using the NEBNext^®^ Ultra™ II DNA Library Prep Kit for Illumina^®^ (New England Biolabs) according to the manufacturer’s instructions with some modifications, and sequencing was performed on an Illumina NovaSeq platform to obtain ∼10 million reads per sample.

### CUT&RUN analysis

Raw sequencing reads were quality-checked using FastQC (v0.12.1). Adapter sequences and low-quality bases were removed using fastp (v0.24.0) with automatic adapter detection for paired-end reads based on per-read overlap analysis. Clean reads were aligned to the mouse reference genome (GRCm38/mm10) using Bowtie2 (v2.5.4) with parameters -I 25 -X 1000 --no-mixed --no-discordant --very-sensitive. Aligned reads were sorted and indexed using SAMtools (v1.21). Mitochondrial reads, unmapped reads, multi-mapped reads, and reads with mapping quality <30 were removed. PCR duplicates were marked using Picard (v3.4.0). Genome-wide coverage tracks (BigWig and BedGraph) were generated using deepTools (v3.5.6) bamCoverage. Peaks were called using MACS2 (v2.2.6; p < 1 × 10□□). Differential binding analysis was performed using MAnorm (v3.16.0), and peaks with MA values >1 or <−1 (p < 0.05) were considered differential. Peaks were annotated to the nearest genes using ChIPseeker (v1.42.0).

### Statistical analysis

Samples were compared using an unpaired, two-tailed Student’s *t*-test, two-way ANOVA, or log-rank (Mantel-Cox) test unless specified otherwise.

## Supporting information

HASHTAG INFORMATION

PEAK ANNOTATION_CONTROL

PEAK ANNOTATION_antiPD-1

## Data and Software Availability

Raw sequence files (accession number: PRJNA1426211) and processed data files for the scRNAseq reported in this paper is deposited in the Gene Expression Omnibus (GEO) database (GEO accession number: GSE320057) and made publicly available at the time of publication. Software used in this study is available online: current version of Cell Ranger: https://support.10xgenomics.com/single-cell-gene-expression/software/downloads/latest; Seurat 4.0: https://satijalab.org/seurat/; ggplot2 3.3.3: https://ggplot2.tidyverse.org/index.html; and ImmGen: https://www.immgen.org.

## Author’s Contributions

Conceptualization, A.S., T.M, A.S.S., and M.M.G.; data curation, A.S., T.M., A.S.S., A.J.S., S.K., and M.M.G.; investigation, A.S., T.M., A.S.S., S.K., N.N.J., and M.M.G.; visualization, A.S., T.M., A.S.S., S.K., M.M.G.; methodology, A.S., T.M., and M.M.G.; data analysis, A.S., T.M., A.S.S., N.N.J., S.S., A.K.J., and K.R.; formal analysis, A.S., N.N.J., K.E.P., K.H.H., B.T.E., K.C., and M.M.G.; validation, A.S. and M.M.G.; writing – original draft, A.S. and M.M.G.; writing – review & editing, A.S., T.M., K.E.P., K.H.H., B.T.E., K.C., A.K.J., and M.M.G.; resources, B.T.E., K.C., and M.M.G.; supervision, M.M.G.; project administration, M.M.G.

## Acknowledgments

M.M.G. was a Cancer Prevention and Research Institute of Texas (CPRIT) Scholar in Cancer Research and an Andrew Sabin Family Foundation Fellow during part of this study. This work was supported by CPRIT (Recruitment of First-Time Tenure-Track Faculty Members; RR190017), an Andrew Sabin Family Foundation Fellowship, a Parker Institute for Cancer Immunotherapy (PICI) Bridge Scholar Award, a University of Texas (UT) Rising Stars Award, and NCI R01CA282027 to M.M.G. and NCI U01CA247760 to K.C. K.H.H. is a CPRIT Scholar in Cancer Research and a PICI and V Foundation Bridge Scholar. K.E.P. was supported by an Andrew Sabin Family Foundation Fellowship at MD Anderson Cancer Center (MDACC), a Melanoma SPORE Developmental Research Program grant at MDACC (grant P50CA221703 from NCI), and the UT Rising Stars Award. AKJ is partially supported by MDACC institutional funds to the Epigenomics Profiling Core. We thank MDACC Epigenomics Profiling Core (EpiCore) for assistance with CUT&RUN assays. The MDACC Flow Cytometry and Cellular Imaging Core Facility and the Advanced Technology Genomics Core (ATGC) are supported by NCI Core grant P30CA016672. We would like to thank David Pollock at ATGC for assistance with scRNA-seq. We would like to thank X. Lily Wang at Baylor College of Medicine MHC Tetramer Core for production of tetramers used in this study. We would like to thank Waikin Chan at MDACC Metabolomics core for assistance with real-time ATP assays.

