## SUPPLEMENTAL FIGURES for "Bhlhe40 Governs T Cell Effector Differentiation with Distinct Requirements in CD4 and CD8 T Cells for Anti-PD-1 and Anti-CTLA-4 Efficacy"

### Slide 1
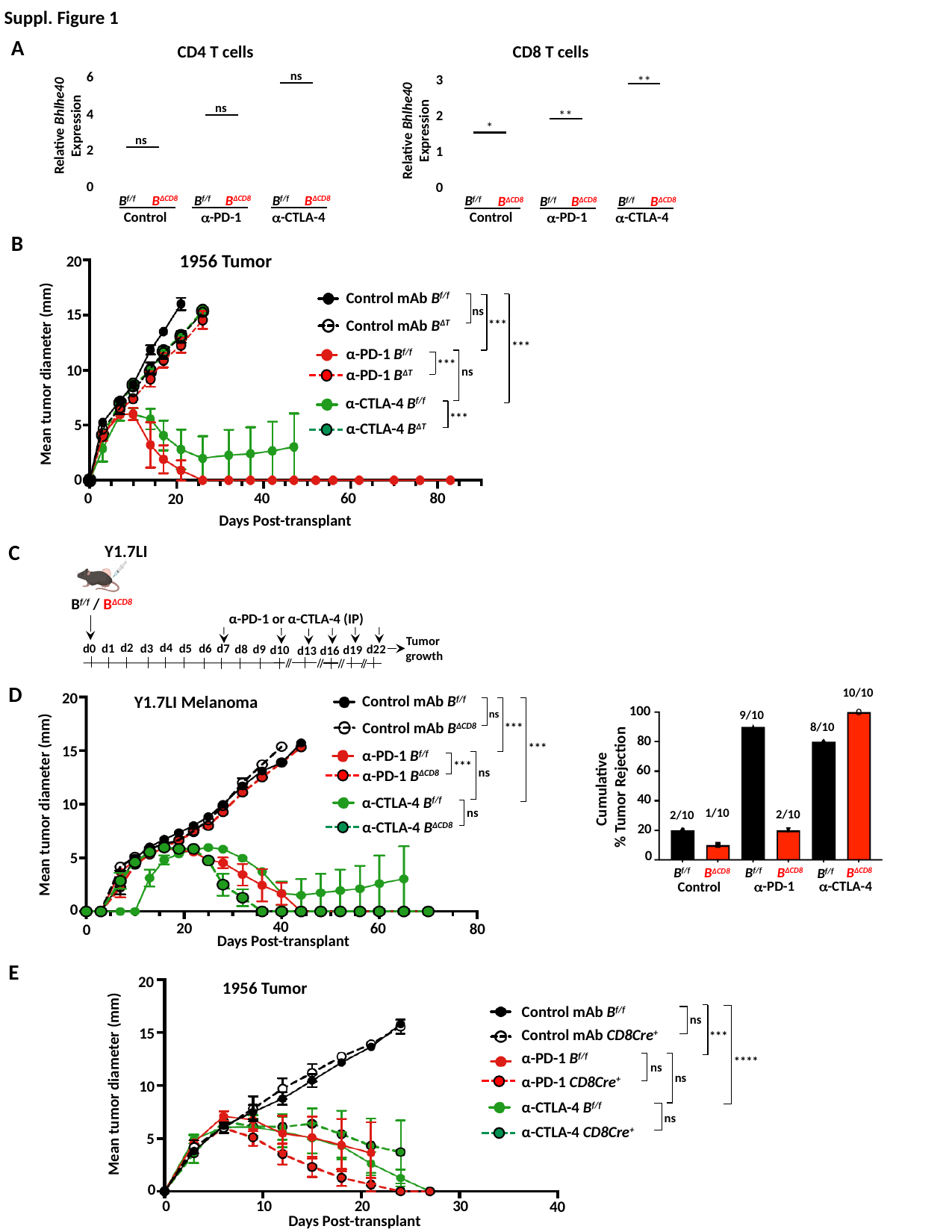

Suppl. Figure 1
A
CD4 T cells
6
ns
ns
Relative Bhlhe40 Expression
4
ns
2
0
B^f/f^
B^ΔCD8^
B^f/f^
B^ΔCD8^
B^f/f^
B^ΔCD8^
Control
α-PD-1
α-CTLA-4
CD8 T cells
3
**
Relative Bhlhe40 Expression
**
2
*
1
0
B^f/f^
B^ΔCD8^
B^f/f^
B^ΔCD8^
B^f/f^
B^ΔCD8^
Control
α-PD-1
α-CTLA-4
B
1956 Tumor
20
Control mAb B^f/f^
ns
Control mAb B^ΔT^
***
***
α-PD-1 B^f/f^
***
ns
α-PD-1 B^ΔT^
α-CTLA-4 B^f/f^
***
α-CTLA-4 B^ΔT^
15
Mean tumor diameter (mm)
10
5
0
80
0
40
60
20
Days Post-transplant
C
Y1.7LI
B^f/f^ / B^ΔCD8^
α-PD-1 or α-CTLA-4 (IP)
d4
 d2
d1
d5
d0
d6
d3
d7
d9
d8
d10
Tumor
growth
d22
d19
d13
d16
10/10
100
80
60
40
20
0
B^f/f^
B^f/f^
B^ΔCD8^
B^ΔCD8^
B^ΔCD8^
B^f/f^
α-CTLA-4
α-PD-1
Control
9/10
8/10
Cumulative
% Tumor Rejection
1/10
2/10
2/10
D
20
Y1.7LI Melanoma
Control mAb B^f/f^
ns
Control mAb B^ΔCD8^
***
***
α-PD-1 B^f/f^
***
ns
α-PD-1 B^ΔCD8^
α-CTLA-4 B^f/f^
ns
α-CTLA-4 B^ΔCD8^
15
Mean tumor diameter (mm)
10
5
0
40
20
60
80
0
Days Post-transplant
E
20
1956 Tumor
Control mAb B^f/f^
Control mAb CD8Cre^+^
α-PD-1 B^f/f^
α-PD-1 CD8Cre^+^
α-CTLA-4 B^f/f^
α-CTLA-4 CD8Cre^+^
ns
15
***
****
ns
Mean tumor diameter (mm)
ns
10
ns
5
0
30
20
10
0
40
Days Post-transplant

### Slide 2
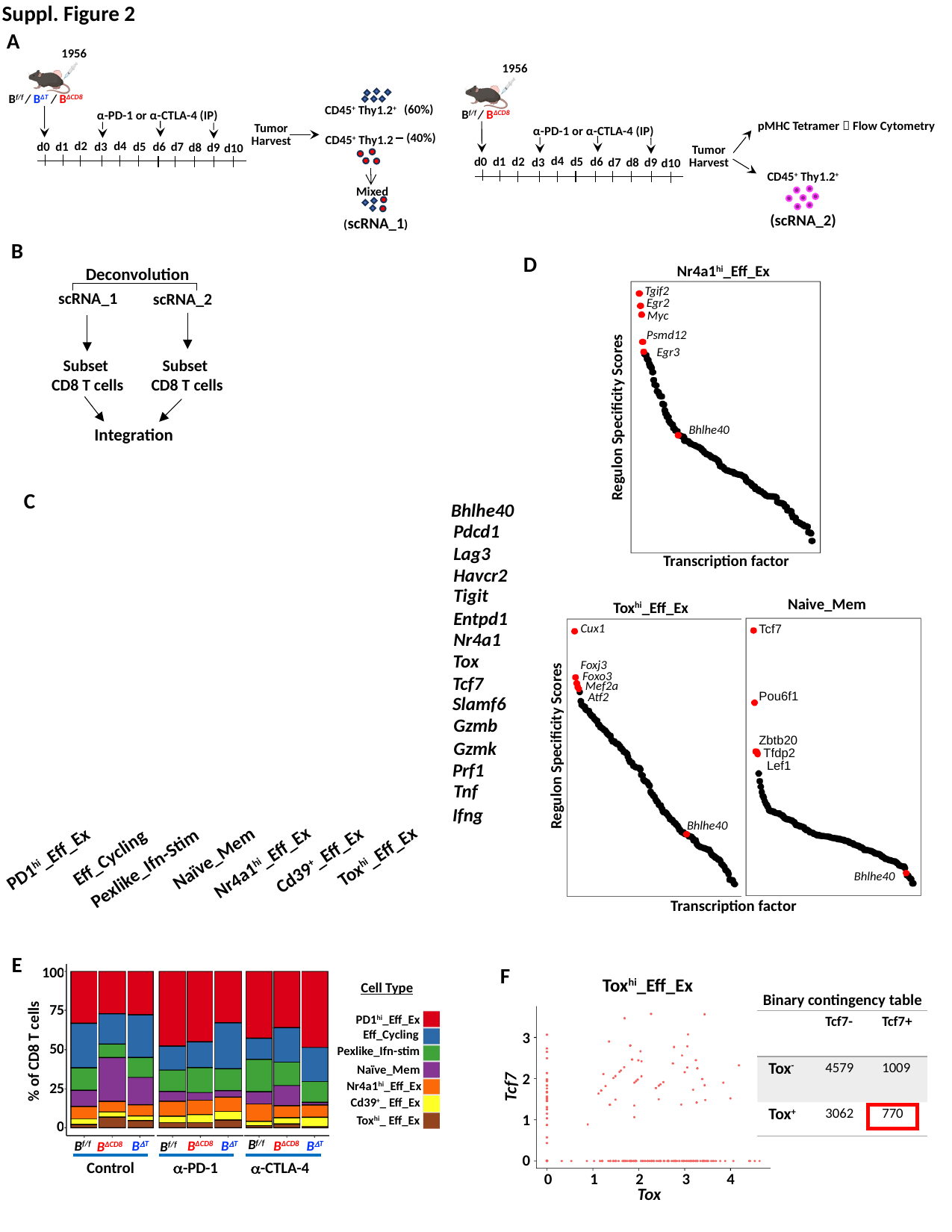

Suppl. Figure 2
A
1956
α-PD-1 or α-CTLA-4 (IP)
d4
 d2
d1
d5
d0
d6
d3
d7
d9
d8
d10
B^f/f^ / B^ΔT^ / B^ΔCD8^
Tumor Harvest
 (60%)
CD45^+^ Thy1.2^+^
CD45^+^ Thy1.2
 (40%)
Mixed
(scRNA_1)
1956
α-PD-1 or α-CTLA-4 (IP)
d4
 d2
d1
d5
d0
d6
d3
d7
d9
d8
d10
B^f/f^ / B^ΔCD8^
pMHC Tetramer 🡪 Flow Cytometry
CD45^+^ Thy1.2^+^
(scRNA_2)
Tumor Harvest
B
Deconvolution
scRNA_1
scRNA_2
Subset
CD8 T cells
Subset
CD8 T cells
Integration
D
Nr4a1^hi^_Eff_Ex
Tgif2
Egr2
Myc
Psmd12
Egr3
Bhlhe40
Regulon Specificity Scores
C
Bhlhe40
Pdcd1
Lag3
Havcr2
Tigit
Gzmb
Gzmk
Prf1
Tnf
Ifng
Entpd1
Nr4a1
Tox
Tcf7
Slamf6
Transcription factor
Naive_Mem
Tcf7
Pou6f1
Zbtb20
Tfdp2
Lef1
Bhlhe40
Tox^hi^_Eff_Ex
Cux1
Foxj3
Foxo3
Mef2a
Atf2
Regulon Specificity Scores
Bhlhe40
Naïve_Mem
Eff_Cycling
Tox^hi^_Eff_Ex
PD1^hi^_Eff_Ex
Cd39^+^_Eff_Ex
Pexlike_Ifn-Stim
Nr4a1^hi^_Eff_Ex
Transcription factor
E
100
75
50
% of CD8 T cells
25
0
Cell Type
Eff_Cycling
Naïve_Mem
PD1^hi^_Eff_Ex
Pexlike_Ifn-stim
Tox^hi^_ Eff_Ex
Cd39^+^_ Eff_Ex
Nr4a1^hi^_Eff_Ex
B^f/f^
B^f/f^
B^ΔCD8^
α-PD-1
Control
α-CTLA-4
B^ΔT^
B^ΔCD8^
B^ΔT^
B^ΔCD8^
B^ΔT^
B^f/f^
F
Tox^hi^_Eff_Ex
Binary contingency table
| | Tcf7- | Tcf7+ |
| --- | --- | --- |
| Tox^-^ | 4579 | 1009 |
| Tox^+^ | 3062 | 770 |
3
2
1
0
Tcf7
0
1
2
3
4
Tox

### Slide 3
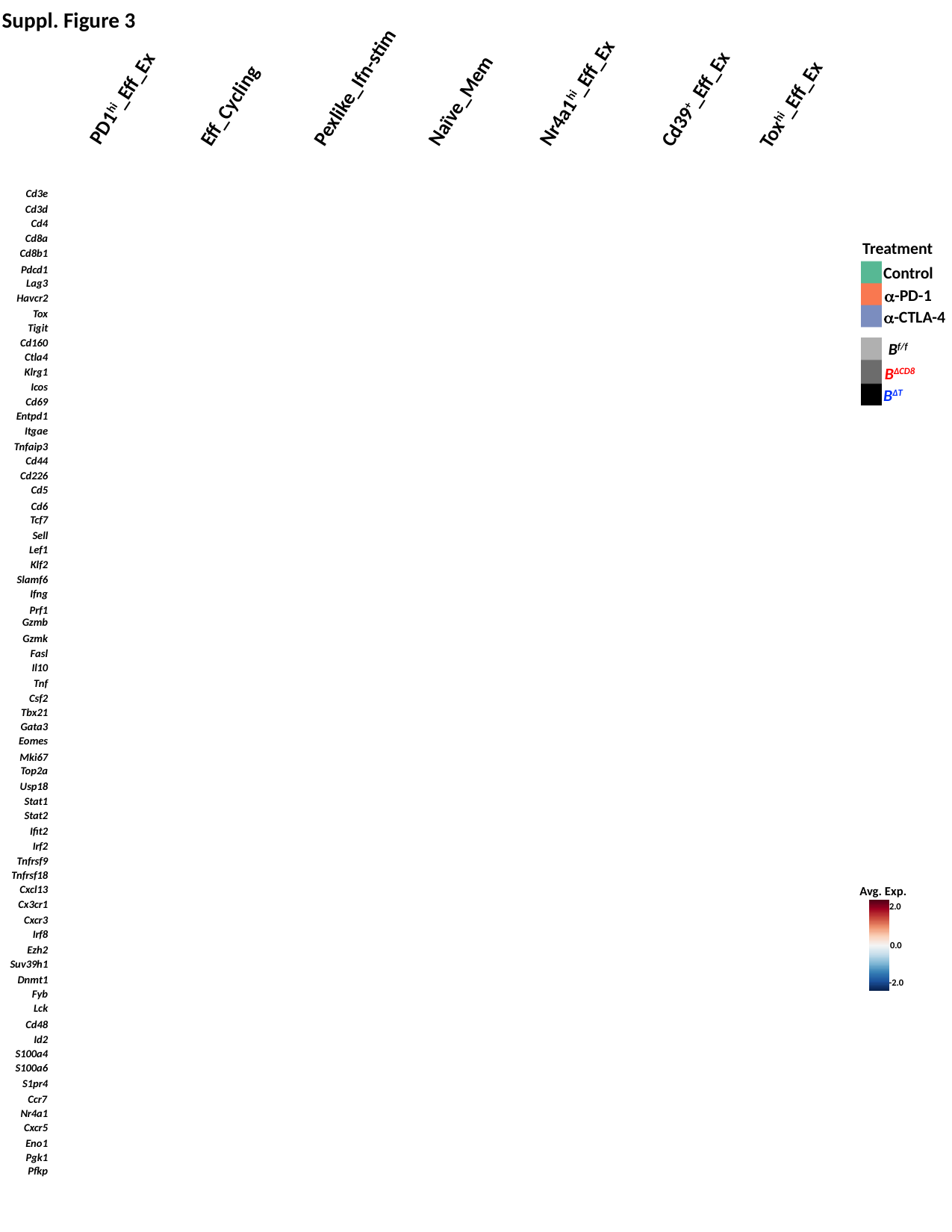

Suppl. Figure 3
PD1^hi^_Eff_Ex
Pexlike_Ifn-stim
Naïve_Mem
Nr4a1^hi^_Eff_Ex
Eff_Cycling
Cd39^+^_Eff_Ex
Tox^hi^_Eff_Ex
Cd3e
Cd3d
Cd4
Cd8a
Cd8b1
Pdcd1
Lag3
Havcr2
Tox
Tigit
Cd160
Ctla4
Klrg1
Icos
Cd69
Entpd1
Itgae
Tnfaip3
Cd44
Cd226
Cd5
Cd6
Tcf7
Sell
Lef1
Klf2
Slamf6
Ifng
Prf1
Gzmb
Gzmk
Fasl
Il10
Tnf
Csf2
Tbx21
Gata3
Eomes
Mki67
Top2a
Usp18
Stat1
Stat2
Ifit2
Irf2
Tnfrsf9
Tnfrsf18
Cxcl13
Cx3cr1
Cxcr3
Irf8
Ezh2
Suv39h1
Dnmt1
Fyb
Lck
Cd48
Id2
S100a4
S100a6
S1pr4
Ccr7
Nr4a1
Cxcr5
Eno1
Pgk1
Pfkp
Treatment
Control
α-PD-1
α-CTLA-4
B^f/f^
B^ΔCD8^
B^ΔT^
Avg. Exp.
2.0
0.0
-2.0

### Slide 4
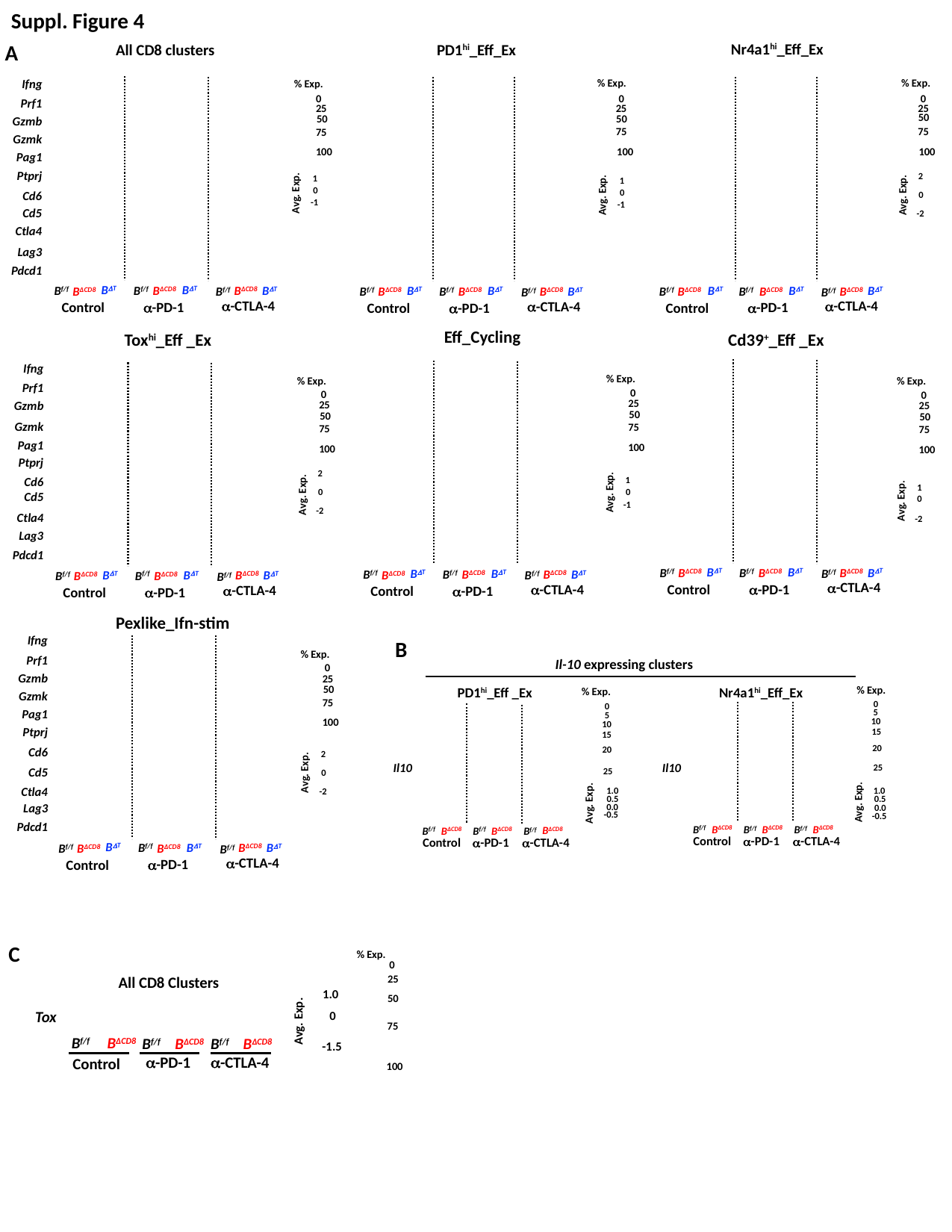

Suppl. Figure 4
A
Nr4a1^hi^_Eff_Ex
All CD8 clusters
PD1^hi^_Eff_Ex
Ifng
Prf1
Gzmb
Gzmk
Pag1
Ptprj
Cd6
Cd5
Ctla4
Lag3
Pdcd1
% Exp.
% Exp.
% Exp.
0
0
0
25
25
25
50
50
50
75
75
75
100
100
100
1
Avg. Exp.
0
-1
2
1
Avg. Exp.
Avg. Exp.
0
0
-1
-2
B^ΔT^
B^f/f^
B^ΔCD8^
α-CTLA-4
α-PD-1
Control
B^ΔT^
B^ΔT^
B^ΔCD8^
B^f/f^
B^ΔCD8^
B^f/f^
B^ΔT^
B^f/f^
B^ΔCD8^
α-CTLA-4
α-PD-1
Control
B^ΔT^
B^ΔT^
B^ΔCD8^
B^f/f^
B^ΔCD8^
B^f/f^
B^ΔT^
B^f/f^
B^ΔCD8^
α-CTLA-4
α-PD-1
Control
B^ΔT^
B^ΔT^
B^ΔCD8^
B^f/f^
B^ΔCD8^
B^f/f^
Eff_Cycling
Cd39^+^_Eff _Ex
Tox^hi^_Eff _Ex
Ifng
Prf1
Gzmb
Gzmk
Pag1
Ptprj
Cd6
Cd5
Ctla4
Lag3
Pdcd1
% Exp.
25
50
75
100
0
% Exp.
25
50
75
100
0
% Exp.
25
50
75
100
0
2
1
Avg. Exp.
1
Avg. Exp.
0
0
Avg. Exp.
0
-1
-2
-2
B^ΔT^
B^f/f^
B^ΔCD8^
α-CTLA-4
α-PD-1
Control
B^ΔT^
B^ΔT^
B^ΔCD8^
B^f/f^
B^ΔCD8^
B^f/f^
B^ΔT^
B^f/f^
B^ΔCD8^
α-CTLA-4
α-PD-1
Control
B^ΔT^
B^ΔT^
B^ΔCD8^
B^f/f^
B^ΔCD8^
B^f/f^
B^ΔT^
B^f/f^
B^ΔCD8^
α-CTLA-4
α-PD-1
Control
B^ΔT^
B^ΔT^
B^ΔCD8^
B^f/f^
B^ΔCD8^
B^f/f^
Pexlike_Ifn-stim
Ifng
Prf1
Gzmb
Gzmk
Pag1
Ptprj
Cd6
Cd5
Ctla4
Lag3
Pdcd1
B
% Exp.
25
50
75
100
0
Il-10 expressing clusters
% Exp.
0
5
10
15
20
25
PD1^hi^_Eff _Ex
Nr4a1^hi^_Eff_Ex
% Exp.
0
5
10
15
20
25
Il10
Il10
1.0
Avg. Exp.
0.5
0.0
-0.5
1.0
Avg. Exp.
0.5
-0.5
0.0
B^ΔCD8^
B^ΔCD8^
B^ΔCD8^
B^f/f^
Control
α-PD-1
α-CTLA-4
B^f/f^
B^f/f^
B^ΔCD8^
B^ΔCD8^
B^ΔCD8^
B^f/f^
Control
α-PD-1
α-CTLA-4
B^f/f^
B^f/f^
2
Avg. Exp.
0
-2
B^ΔT^
B^f/f^
B^ΔCD8^
α-CTLA-4
α-PD-1
Control
B^ΔT^
B^ΔT^
B^ΔCD8^
B^f/f^
B^ΔCD8^
B^f/f^
C
1.0
Avg. Exp.
0
-1.5
% Exp.
25
50
75
100
0
All CD8 Clusters
Tox
B^f/f^
B^ΔCD8^
B^ΔCD8^
B^f/f^
B^ΔCD8^
B^f/f^
α-CTLA-4
α-PD-1
Control

### Slide 5
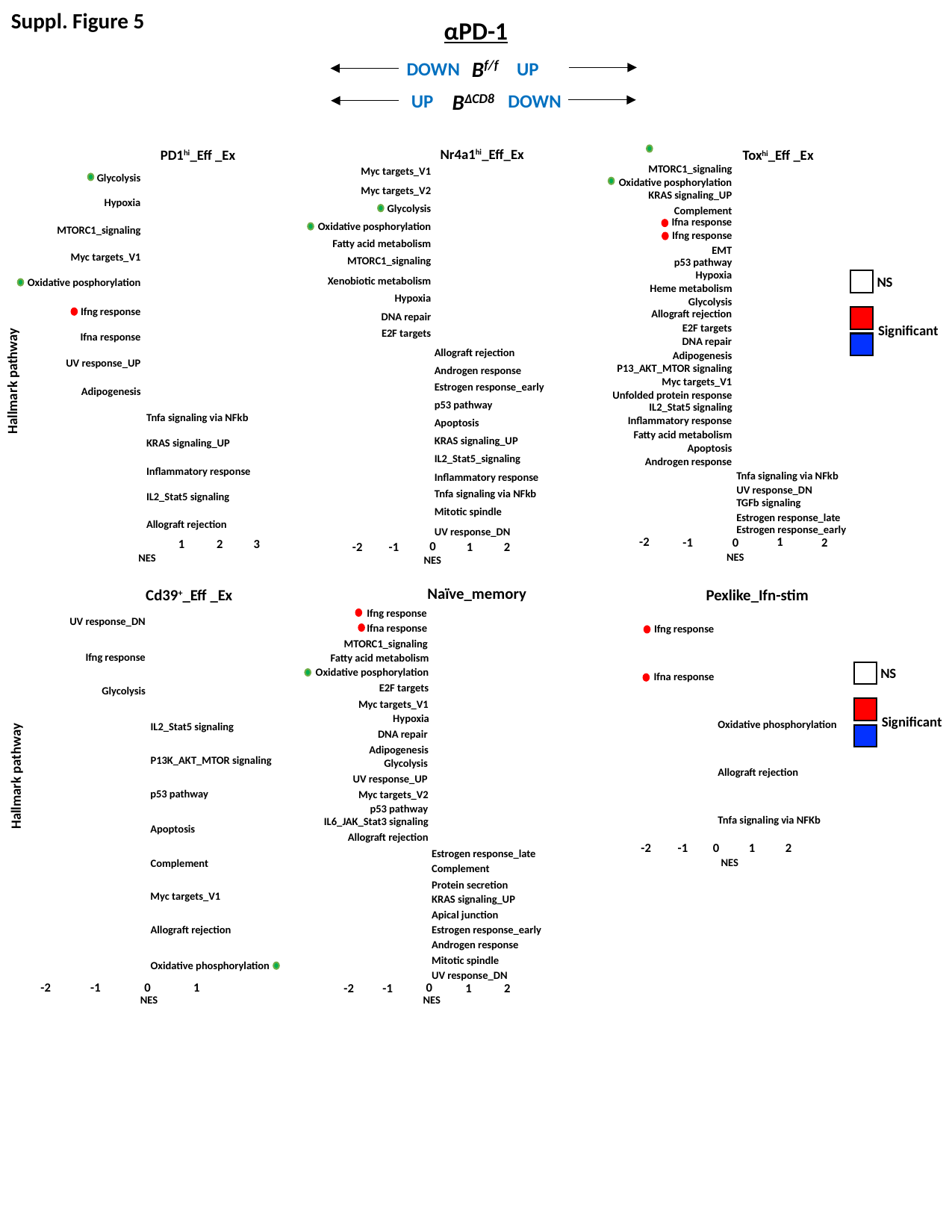

Suppl. Figure 5
αPD-1
B^f/f^
DOWN
UP
B^ΔCD8^
UP
DOWN
Nr4a1^hi^_Eff_Ex
PD1^hi^_Eff _Ex
Tox^hi^_Eff _Ex
MTORC1_signaling
Oxidative posphorylation
KRAS signaling_UP
Complement
Ifna response
Ifng response
EMT
p53 pathway
Hypoxia
Glycolysis
Allograft rejection
E2F targets
DNA repair
Adipogenesis
Myc targets_V1
Inflammatory response
Apoptosis
Androgen response
Heme metabolism
P13_AKT_MTOR signaling
Unfolded protein response
IL2_Stat5 signaling
Fatty acid metabolism
Myc targets_V1
Myc targets_V2
Glycolysis
Oxidative posphorylation
Fatty acid metabolism
MTORC1_signaling
Xenobiotic metabolism
Hypoxia
DNA repair
E2F targets
Glycolysis
Hypoxia
MTORC1_signaling
Myc targets_V1
Oxidative posphorylation
Ifng response
Ifna response
UV response_UP
Adipogenesis
NS
Significant
Allograft rejection
Androgen response
Estrogen response_early
p53 pathway
Apoptosis
KRAS signaling_UP
IL2_Stat5_signaling
Inflammatory response
Tnfa signaling via NFkb
Mitotic spindle
UV response_DN
Hallmark pathway
Tnfa signaling via NFkb
KRAS signaling_UP
Inflammatory response
IL2_Stat5 signaling
Allograft rejection
Tnfa signaling via NFkb
UV response_DN
TGFb signaling
Estrogen response_late
Estrogen response_early
-2
1
-1
0
2
1
2
3
0
1
-1
-2
2
NES
NES
NES
Naïve_memory
Pexlike_Ifn-stim
Cd39^+^_Eff _Ex
Ifng response
Ifna response
MTORC1_signaling
Oxidative posphorylation
E2F targets
Myc targets_V1
Hypoxia
DNA repair
Adipogenesis
Glycolysis
p53 pathway
Fatty acid metabolism
IL6_JAK_Stat3 signaling
UV response_UP
Myc targets_V2
Allograft rejection
UV response_DN
Ifng response
Ifna response
Ifng response
NS
Significant
Glycolysis
Oxidative phosphorylation
Allograft rejection
Tnfa signaling via NFKb
IL2_Stat5 signaling
P13K_AKT_MTOR signaling
p53 pathway
Apoptosis
Complement
Myc targets_V1
Allograft rejection
Oxidative phosphorylation
Hallmark pathway
-2
0
1
2
-1
Estrogen response_late
Complement
Protein secretion
KRAS signaling_UP
Apical junction
Estrogen response_early
Androgen response
Mitotic spindle
UV response_DN
NES
-1
0
1
0
-2
-2
-1
1
2
NES
NES

### Slide 6
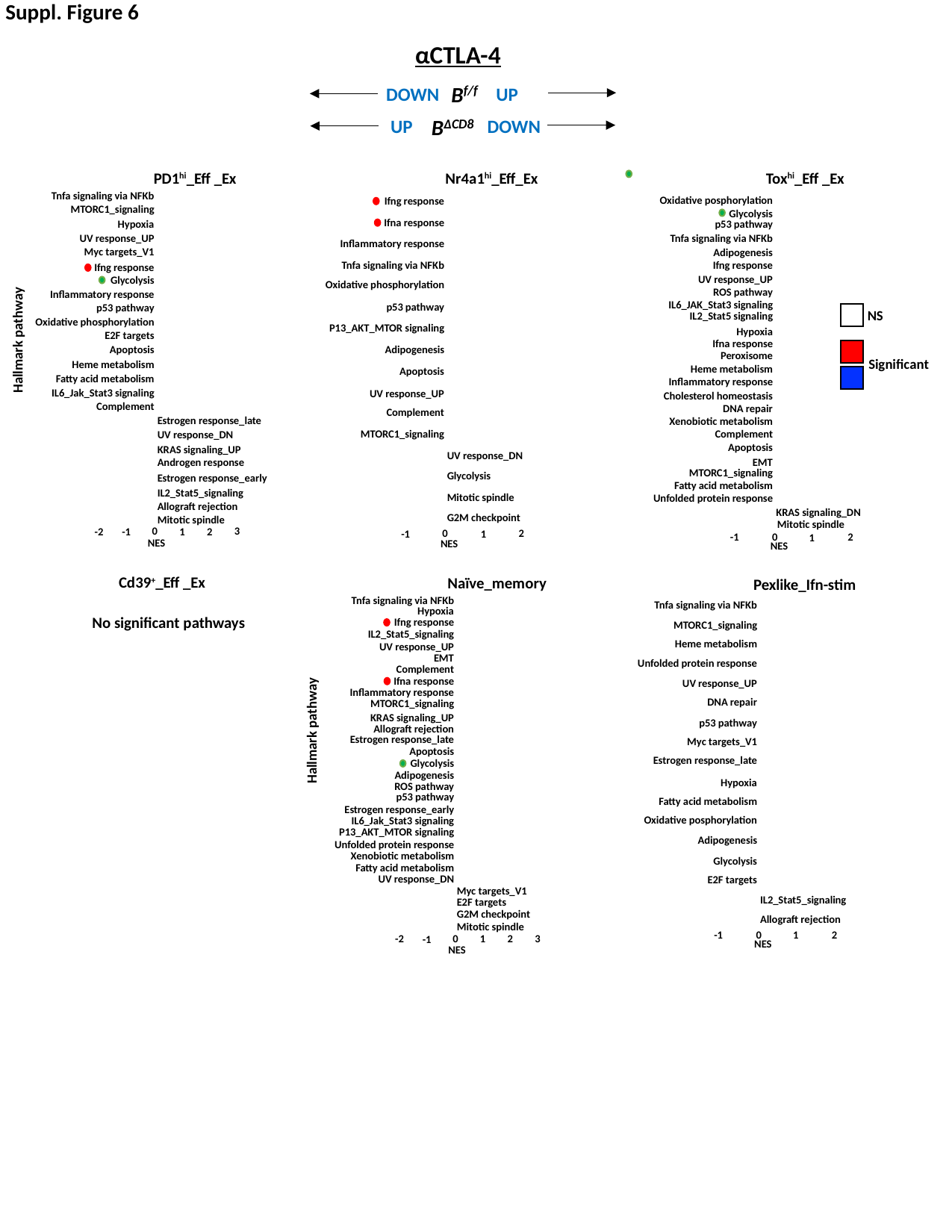

Suppl. Figure 6
αCTLA-4
B^f/f^
DOWN
UP
B^ΔCD8^
UP
DOWN
PD1^hi^_Eff _Ex
Nr4a1^hi^_Eff_Ex
Tox^hi^_Eff _Ex
Tnfa signaling via NFKb
MTORC1_signaling
Hypoxia
UV response_UP
Myc targets_V1
Ifng response
Glycolysis
Inflammatory response
p53 pathway
Oxidative phosphorylation
E2F targets
Apoptosis
Heme metabolism
Fatty acid metabolism
IL6_Jak_Stat3 signaling
Complement
Oxidative posphorylation
Glycolysis
p53 pathway
Tnfa signaling via NFKb
Adipogenesis
Ifng response
UV response_UP
ROS pathway
IL6_JAK_Stat3 signaling
IL2_Stat5 signaling
Hypoxia
Ifna response
Peroxisome
Heme metabolism
Inflammatory response
Cholesterol homeostasis
DNA repair
Xenobiotic metabolism
Complement
Apoptosis
EMT
MTORC1_signaling
Fatty acid metabolism
Unfolded protein response
Ifng response
Ifna response
Inflammatory response
Tnfa signaling via NFKb
Oxidative phosphorylation
p53 pathway
NS
Significant
Hallmark pathway
P13_AKT_MTOR signaling
Adipogenesis
Apoptosis
UV response_UP
Complement
Estrogen response_late
UV response_DN
KRAS signaling_UP
Androgen response
Estrogen response_early
IL2_Stat5_signaling
Allograft rejection
Mitotic spindle
MTORC1_signaling
UV response_DN
Glycolysis
Mitotic spindle
KRAS signaling_DN
G2M checkpoint
Mitotic spindle
3
0
1
2
-2
-1
0
2
-1
1
2
-1
0
1
NES
NES
NES
Cd39^+^_Eff _Ex
Naïve_memory
Pexlike_Ifn-stim
Tnfa signaling via NFKb
Hypoxia
Ifng response
IL2_Stat5_signaling
UV response_UP
EMT
Complement
Ifna response
Inflammatory response
MTORC1_signaling
KRAS signaling_UP
Allograft rejection
Estrogen response_late
Apoptosis
Glycolysis
Adipogenesis
ROS pathway
p53 pathway
Estrogen response_early
IL6_Jak_Stat3 signaling
P13_AKT_MTOR signaling
Unfolded protein response
Xenobiotic metabolism
Fatty acid metabolism
UV response_DN
Tnfa signaling via NFKb
MTORC1_signaling
Heme metabolism
Unfolded protein response
UV response_UP
DNA repair
p53 pathway
Myc targets_V1
Estrogen response_late
Hypoxia
Fatty acid metabolism
Oxidative posphorylation
Adipogenesis
Glycolysis
E2F targets
No significant pathways
Hallmark pathway
Myc targets_V1
E2F targets
G2M checkpoint
Mitotic spindle
IL2_Stat5_signaling
Allograft rejection
2
-1
0
1
3
0
1
2
-2
-1
NES
NES

### Slide 7
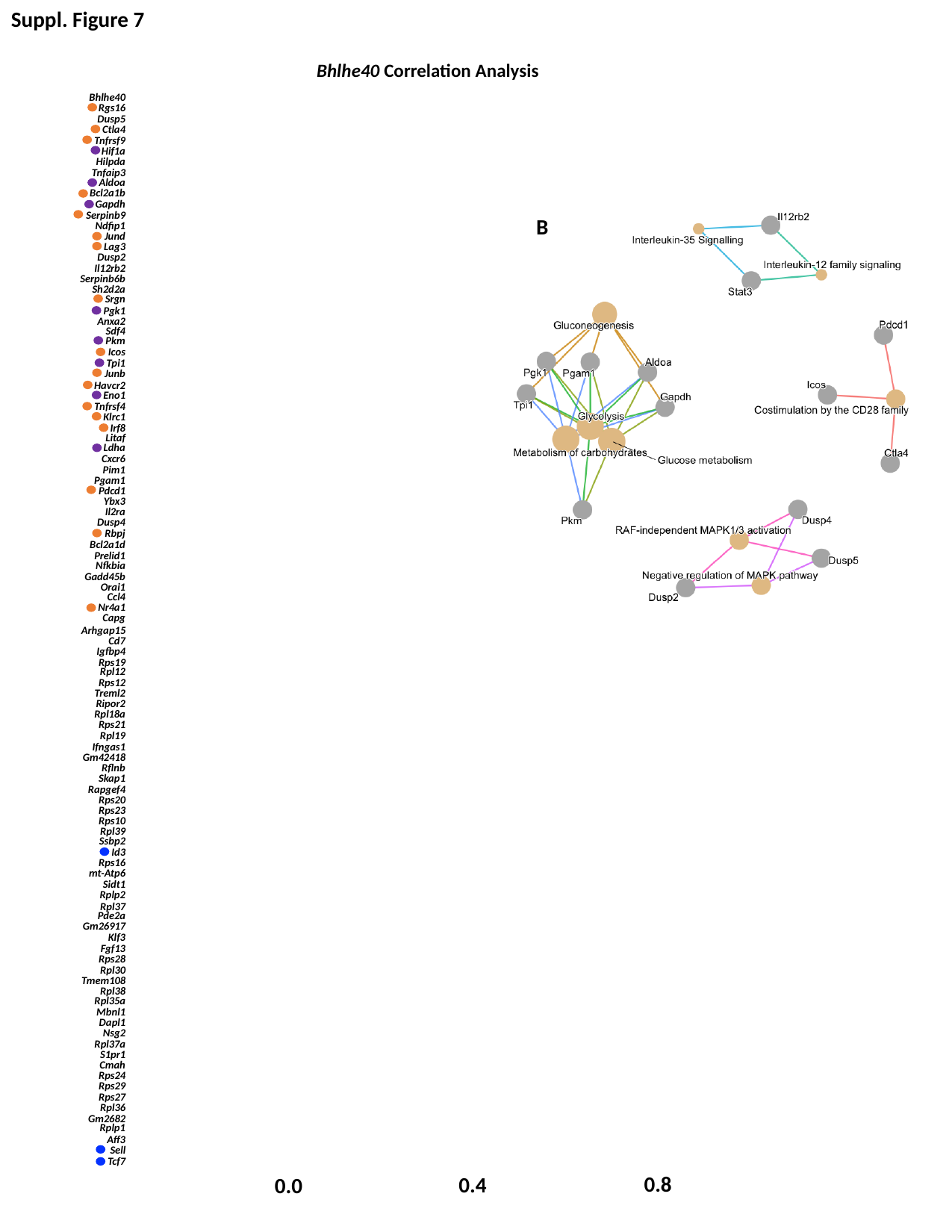

Suppl. Figure 7
Bhlhe40 Correlation Analysis
Bhlhe40
Rgs16
Dusp5
Ctla4
Tnfrsf9
Hif1a
Hilpda
Tnfaip3
Aldoa
Bcl2a1b
Gapdh
Serpinb9
Ndfip1
Jund
Lag3
Dusp2
Il12rb2
Serpinb6b
Sh2d2a
Srgn
Pgk1
Anxa2
Sdf4
Pkm
Icos
Tpi1
Junb
Havcr2
Eno1
Tnfrsf4
Klrc1
Irf8
Litaf
Ldha
Cxcr6
Pim1
Pgam1
Pdcd1
Ybx3
Il2ra
Dusp4
Rbpj
Bcl2a1d
Prelid1
Nfkbia
Gadd45b
Orai1
Ccl4
Nr4a1
Capg
Arhgap15
Cd7
Igfbp4
Rps19
Rpl12
Rps12
Treml2
Ripor2
Rpl18a
Rps21
Rpl19
Ifngas1
Gm42418
Rflnb
Skap1
Rapgef4
Rps20
Rps23
Rps10
Rpl39
Ssbp2
Id3
Rps16
mt-Atp6
Sidt1
Rplp2
Rpl37
Pde2a
Gm26917
Klf3
Fgf13
Rps28
Rpl30
Tmem108
Rpl38
Rpl35a
Mbnl1
Dapl1
Nsg2
Rpl37a
S1pr1
Cmah
Rps24
Rps29
Rps27
Rpl36
Gm2682
Rplp1
Aff3
Sell
Tcf7
B
0.8
0.4
0.0

### Slide 8
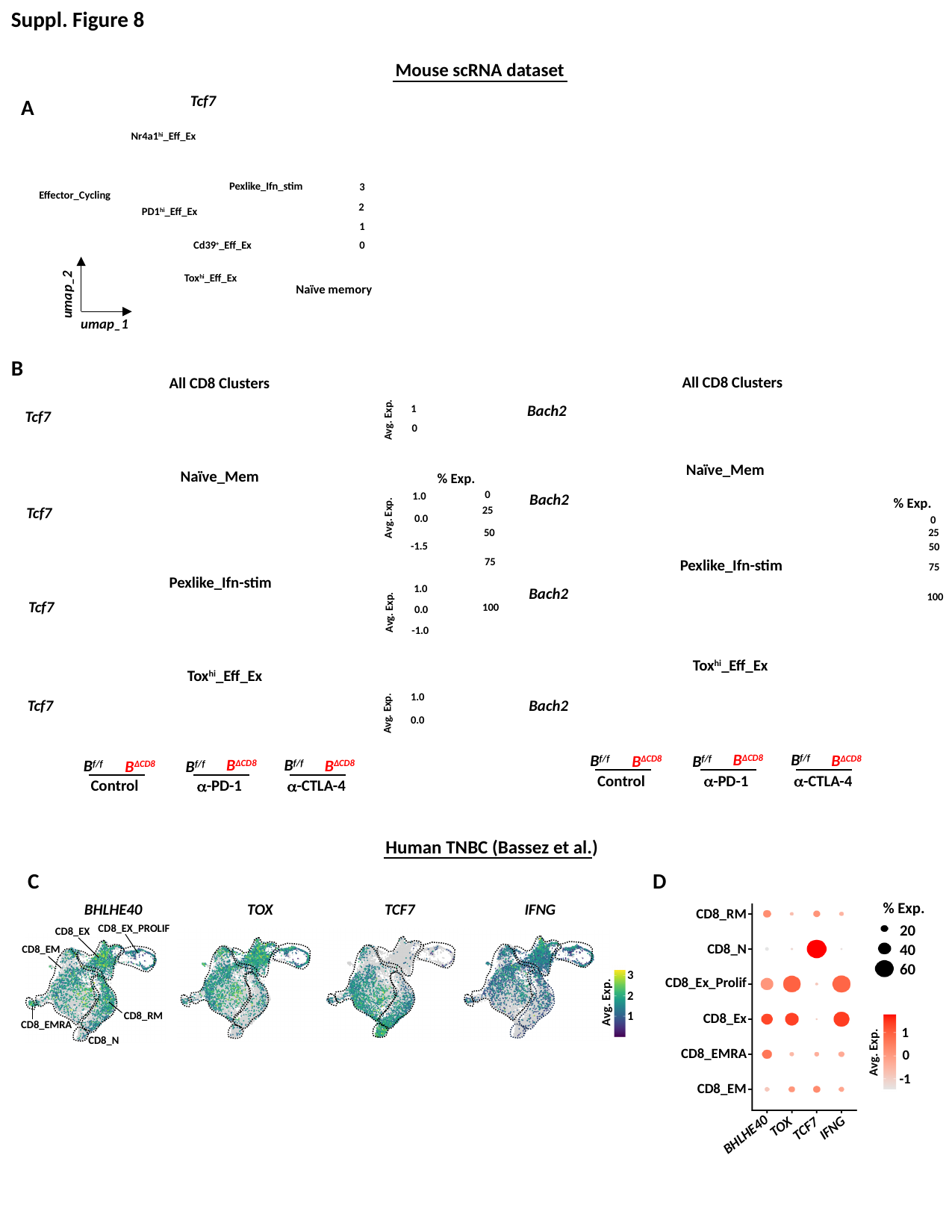

Suppl. Figure 8
Mouse scRNA dataset
Tcf7
Nr4a1^hi^_Eff_Ex
Pexlike_Ifn_stim
3
2
1
0
Effector_Cycling
PD1^hi^_Eff_Ex
Cd39^+^_Eff_Ex
umap_2
Tox^hi^_Eff_Ex
Naïve memory
umap_1
A
B
All CD8 Clusters
Bach2
Naïve_Mem
Bach2
Pexlike_Ifn-stim
Bach2
Tox^hi^_Eff_Ex
Bach2
B^ΔCD8^
B^f/f^
B^ΔCD8^
B^f/f^
B^ΔCD8^
B^f/f^
Control
α-PD-1
α-CTLA-4
All CD8 Clusters
1
Avg. Exp.
0
Tcf7
Naïve_Mem
Tcf7
% Exp.
0
25
50
75
100
1.0
% Exp.
0
25
50
75
100
Avg. Exp.
0.0
-1.5
Pexlike_Ifn-stim
Tcf7
1.0
Avg. Exp.
0.0
-1.0
Tox^hi^_Eff_Ex
Tcf7
1.0
Avg. Exp.
0.0
B^ΔCD8^
B^f/f^
B^ΔCD8^
B^f/f^
B^ΔCD8^
B^f/f^
Control
α-PD-1
α-CTLA-4
Human TNBC (Bassez et al.)
BHLHE40
TOX
TCF7
IFNG
CD8_EX_PROLIF
CD8_EX
CD8_EM
CD8_RM
CD8_EMRA
CD8_N
3
2
1
Avg. Exp.
C
D
% Exp.
20
40
60
1
Avg. Exp.
0
-1
CD8_RM
CD8_N
CD8_Ex_Prolif
CD8_Ex
CD8_EMRA
CD8_EM
BHLHE40
TOX
IFNG
TCF7

### Slide 9
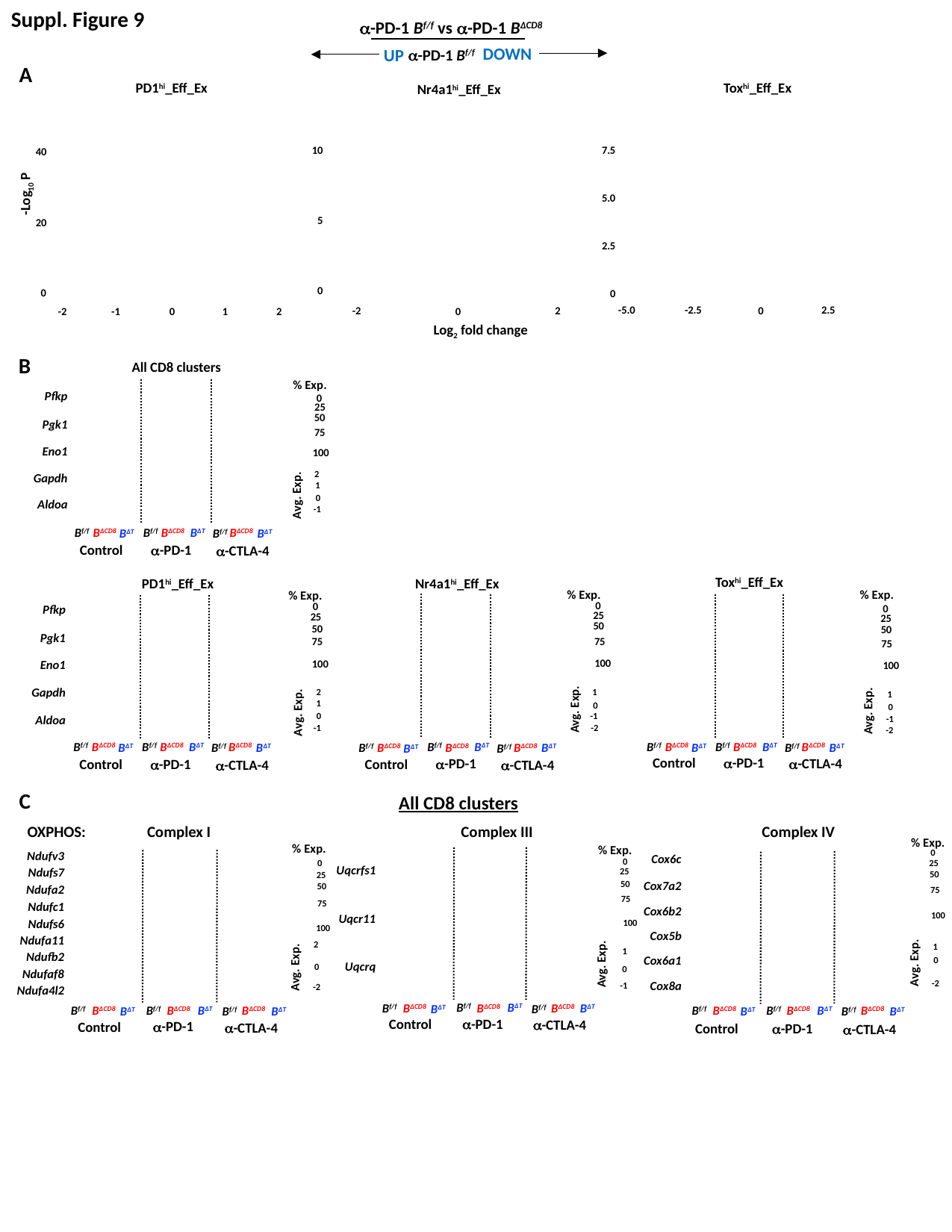

Suppl. Figure 9
α-PD-1 B^f/f^ vs α-PD-1 B^ΔCD8^
DOWN
UP
α-PD-1 B^f/f^
A
Tox^hi^_Eff_Ex
PD1^hi^_Eff_Ex
Nr4a1^hi^_Eff_Ex
7.5
10
40
-Log_10_ P
5.0
5
20
2.5
0
0
0
-5.0
-2.5
2.5
0
2
-2
0
-1
1
2
-2
0
Log_2_ fold change
B
All CD8 clusters
% Exp.
Pfkp
Pgk1
Eno1
Gapdh
Aldoa
0
25
50
75
100
2
1
Avg. Exp.
0
-1
B^ΔT^
B^f/f^
B^f/f^
B^ΔCD8^
α-PD-1
Control
α-CTLA-4
B^ΔCD8^
B^ΔCD8^
B^f/f^
B^ΔT^
B^ΔT^
Tox^hi^_Eff_Ex
PD1^hi^_Eff_Ex
Nr4a1^hi^_Eff_Ex
% Exp.
0
25
50
75
100
1
Avg. Exp.
0
-1
-2
% Exp.
% Exp.
0
0
Pfkp
Pgk1
Eno1
Gapdh
Aldoa
25
25
50
50
75
75
100
100
2
1
1
0
Avg. Exp.
Avg. Exp.
0
-1
-1
-2
B^ΔT^
B^f/f^
B^f/f^
B^ΔCD8^
α-PD-1
Control
α-CTLA-4
B^ΔCD8^
B^ΔCD8^
B^f/f^
B^ΔT^
B^ΔT^
B^ΔT^
B^f/f^
B^f/f^
B^ΔCD8^
α-PD-1
Control
α-CTLA-4
B^ΔCD8^
B^ΔCD8^
B^f/f^
B^ΔT^
B^ΔT^
B^ΔT^
B^f/f^
B^f/f^
B^ΔCD8^
α-PD-1
Control
α-CTLA-4
B^ΔCD8^
B^ΔCD8^
B^f/f^
B^ΔT^
B^ΔT^
C
All CD8 clusters
OXPHOS:
Complex I
Complex III
Complex IV
% Exp.
0
25
50
75
100
1
Avg. Exp.
0
-2
% Exp.
0
25
50
75
100
2
Avg. Exp.
0
-2
% Exp.
0
25
50
75
100
Ndufv3
Ndufs7
Ndufa2
Ndufc1
Ndufs6
Ndufa11
Ndufb2
Ndufaf8
Ndufa4l2
Cox6c
Cox7a2
Cox6b2
Cox5b
Cox6a1
Cox8a
Uqcrfs1
Uqcr11
Uqcrq
1
Avg. Exp.
0
-1
B^ΔT^
B^f/f^
B^f/f^
B^ΔCD8^
α-PD-1
Control
α-CTLA-4
B^ΔCD8^
B^ΔCD8^
B^f/f^
B^ΔT^
B^ΔT^
B^ΔT^
B^f/f^
B^f/f^
B^ΔCD8^
α-PD-1
Control
α-CTLA-4
B^ΔCD8^
B^ΔCD8^
B^f/f^
B^ΔT^
B^ΔT^
B^ΔT^
B^f/f^
B^f/f^
B^ΔCD8^
α-PD-1
Control
α-CTLA-4
B^ΔCD8^
B^ΔCD8^
B^f/f^
B^ΔT^
B^ΔT^

### Slide 10
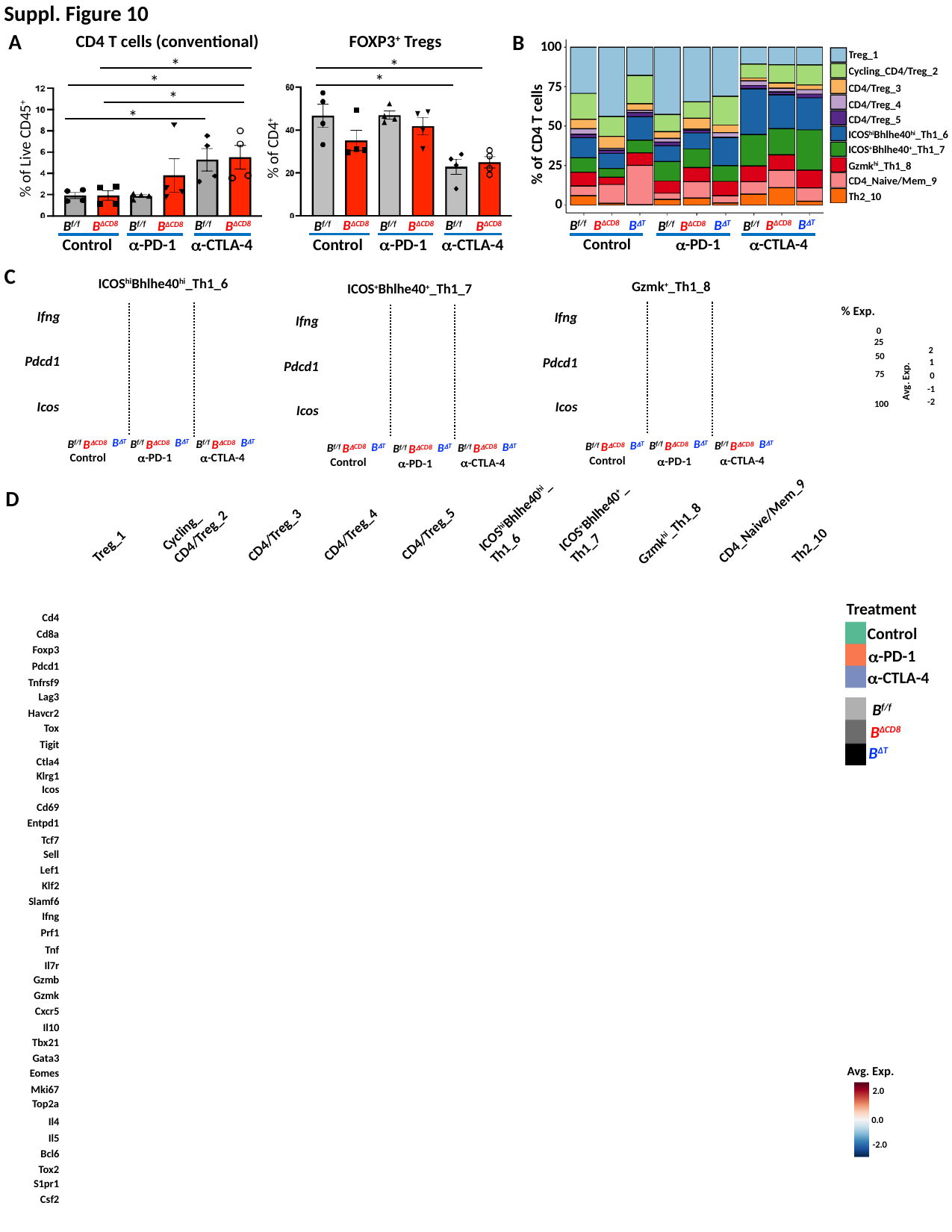

Suppl. Figure 10
A
CD4 T cells (conventional)
*
*
*
*
% of Live CD45^+^
B^f/f^
B^ΔCD8^
B^f/f^
B^ΔCD8^
B^ΔCD8^
α-PD-1
α-CTLA-4
Control
B^f/f^
FOXP3^+^ Tregs
*
*
% of CD4^+^
B^f/f^
B^ΔCD8^
B^f/f^
B^ΔCD8^
B^ΔCD8^
α-PD-1
α-CTLA-4
Control
B^f/f^
B
100
Treg_1
Cycling_CD4/Treg_2
CD4/Treg_3
CD4/Treg_4
CD4/Treg_5
ICOS^hi^Bhlhe40^hi^_Th1_6
ICOS^+^Bhlhe40^+^_Th1_7
Gzmk^hi^_Th1_8
CD4_Naive/Mem_9
Th2_10
75
50
% of CD4 T cells
25
0
B^ΔT^
B^ΔCD8^
B^f/f^
Control
α-CTLA-4
α-PD-1
B^ΔT^
B^ΔT^
B^ΔCD8^
B^ΔCD8^
B^f/f^
B^f/f^
C
ICOS^hi^Bhlhe40^hi^_Th1_6
Ifng
Pdcd1
Icos
B^ΔT^
B^f/f^
B^ΔCD8^
B^ΔT^
B^f/f^
B^ΔCD8^
B^ΔT^
B^f/f^
B^ΔCD8^
Control
α-PD-1
α-CTLA-4
Gzmk^+^_Th1_8
B^ΔT^
B^f/f^
B^ΔCD8^
B^ΔT^
B^f/f^
B^ΔCD8^
B^ΔT^
B^f/f^
B^ΔCD8^
Control
α-CTLA-4
α-PD-1
Ifng
Pdcd1
Icos
ICOS^+^Bhlhe40^+^_Th1_7
B^ΔT^
B^f/f^
B^ΔCD8^
B^ΔT^
B^f/f^
B^ΔCD8^
B^ΔT^
B^f/f^
B^ΔCD8^
Control
α-CTLA-4
α-PD-1
Ifng
Pdcd1
Icos
% Exp.
0
2
1
Avg. Exp.
0
-1
-2
25
50
75
100
ICOS^hi^Bhlhe40^hi^_
Th1_6
D
ICOS^+^Bhlhe40^+^_
Th1_7
Cycling_
CD4/Treg_2
CD4_Naive/Mem_9
Gzmk^hi^_Th1_8
CD4/Treg_5
Treg_1
CD4/Treg_4
CD4/Treg_3
Th2_10
Treatment
Cd4
Cd8a
Foxp3
Pdcd1
Tnfrsf9
Lag3
Havcr2
Tox
Tigit
Ctla4
Klrg1
Icos
Cd69
Entpd1
Tcf7
Sell
Lef1
Klf2
Slamf6
Ifng
Prf1
Tnf
Il7r
Gzmb
Gzmk
Cxcr5
Il10
Tbx21
Gata3
Eomes
Mki67
Top2a
Il4
Il5
Bcl6
Tox2
S1pr1
Csf2
Control
α-PD-1
α-CTLA-4
B^f/f^
B^ΔCD8^
B^ΔT^
Avg. Exp.
2.0
0.0
-2.0

### Slide 11
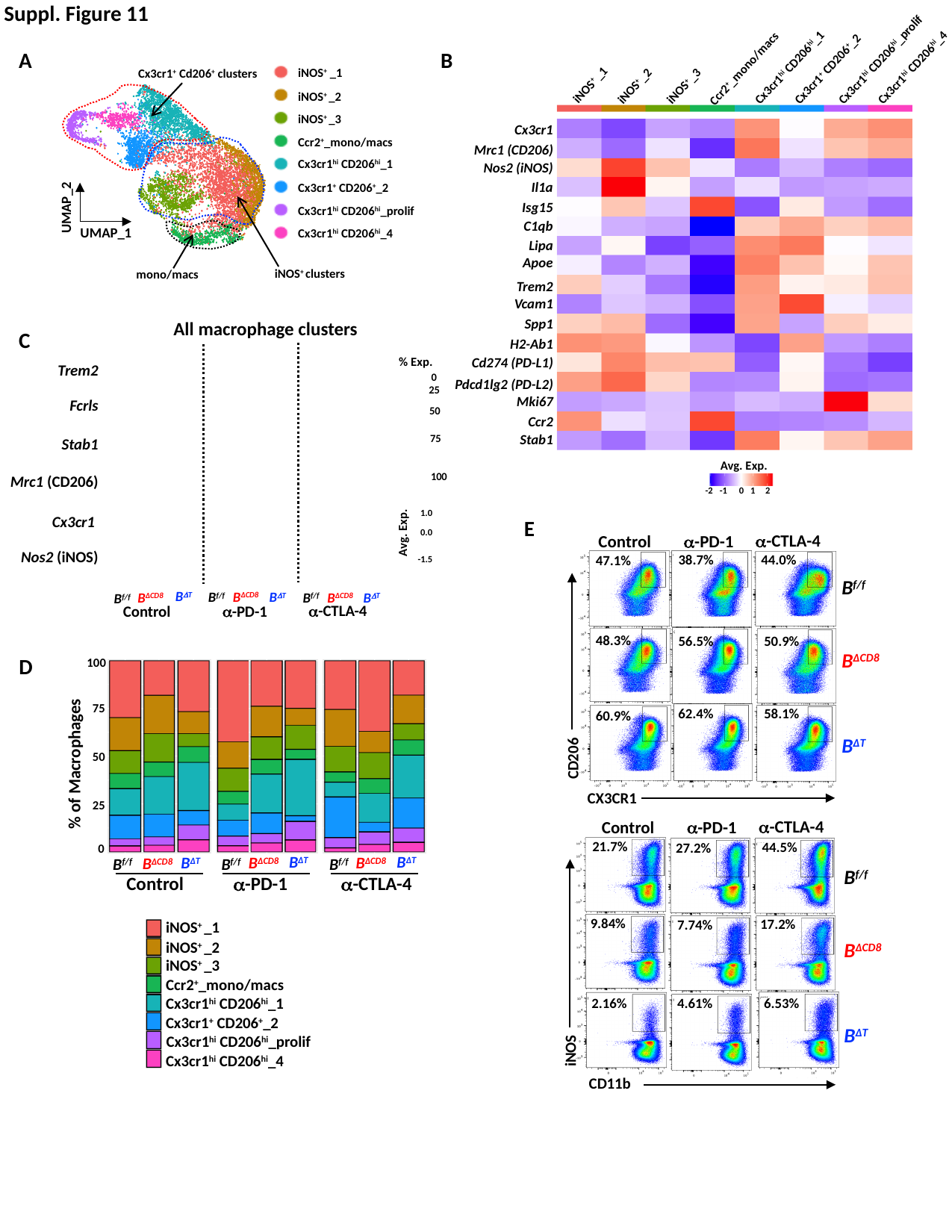

Cx3cr1^hi^ CD206^hi^_prolif
Cx3cr1^+^ CD206^+^_2
Cx3cr1^hi^ CD206^hi^_4
B
Cx3cr1^hi^ CD206^hi^_1
Ccr2^+^_mono/macs
iNOS^+^ _1
iNOS^+^ _2
iNOS^+^ _3
Cx3cr1
Mrc1 (CD206)
Nos2 (iNOS)
Il1a
Isg15
C1qb
Lipa
Apoe
Trem2
Vcam1
Spp1
H2-Ab1
Cd274 (PD-L1)
Pdcd1lg2 (PD-L2)
Mki67
Ccr2
Stab1
Avg. Exp.
-2
-1
0
1
2
Suppl. Figure 11
A
iNOS^+^ _1
iNOS^+^ _2
iNOS^+^ _3
Ccr2^+^_mono/macs
Cx3cr1^hi^ CD206^hi^_1
Cx3cr1^+^ CD206^+^_2
Cx3cr1^hi^ CD206^hi^_prolif
Cx3cr1^hi^ CD206^hi^_4
Cx3cr1^+^ Cd206^+^ clusters
UMAP_2
UMAP_1
iNOS^+^ clusters
mono/macs
All macrophage clusters
C
% Exp.
Trem2
0
25
Fcrls
50
75
Stab1
100
Mrc1 (CD206)
1.0
Cx3cr1
E
α-CTLA-4
α-PD-1
Control
38.7%
44.0%
47.1%
B^f/f^
B^ΔCD8^
B^ΔT^
CD206
CX3CR1
48.3%
50.9%
56.5%
62.4%
58.1%
60.9%
α-CTLA-4
α-PD-1
Control
21.7%
44.5%
27.2%
9.84%
17.2%
7.74%
6.53%
2.16%
4.61%
iNOS
CD11b
B^f/f^
B^ΔCD8^
B^ΔT^
Avg. Exp.
0.0
Nos2 (iNOS)
-1.5
B^ΔT^
B^ΔCD8^
B^f/f^
α-CTLA-4
α-PD-1
Control
B^ΔCD8^
B^ΔT^
B^f/f^
B^ΔCD8^
B^ΔT^
B^f/f^
D
100
75
50
% of Macrophages
25
0
B^ΔT^
B^ΔCD8^
B^f/f^
B^ΔT^
B^ΔCD8^
B^f/f^
B^ΔT^
B^ΔCD8^
B^f/f^
Control
α-PD-1
α-CTLA-4
iNOS^+^ _1
iNOS^+^ _2
iNOS^+^ _3
Ccr2^+^_mono/macs
Cx3cr1^hi^ CD206^hi^_1
Cx3cr1^+^ CD206^+^_2
Cx3cr1^hi^ CD206^hi^_4
Cx3cr1^hi^ CD206^hi^_prolif
